# MAPK/ERK Activation in Macrophages Promotes *Leishmania* Internalization and Pathogenesis

**DOI:** 10.1101/2022.12.08.517760

**Authors:** Umaru Barrie, Katherine Floyd, Arani Datta, Dawn M. Wetzel

## Abstract

The obligate intracellular protozoan parasite *Leishmania* binds several host cell receptors to trigger its uptake by phagocytic cells, ultimately resulting in visceral or cutaneous leishmaniasis. After *Leishmania* engages receptors on macrophages and other phagocytes, a series of signaling pathways in the host cell are activated during its internalization, which are critical for establishment and persistence of *Leishmania* infection. Thus, preventing *Leishmania* internalization by phagocytes could be a novel therapeutic strategy for leishmaniasis. However, the host cellular machinery that mediates promastigote and amastigote uptake is not well understood. Here, using small molecule inhibitors of Mitogen-activated protein kinases/Extracellular signal regulated kinases (MAPK/ERK), we demonstrate that ERK1/2 mediates *Leishmania amazonensis* uptake and (to a lesser extent) phagocytosis of beads by macrophages. We find that inhibition of MEK1/2 or ERK1/2 leads to inefficient amastigote uptake by macrophages. Moreover, using inhibitors and primary macrophages lacking spleen tyrosine kinase (SYK) or Abl family kinases, we show that SYK and Abl family kinases mediate Raf, MEK, and ERK1/2 activity and are necessary for efficient uptake. Finally, we demonstrate that trametinib, a MEK1/2 inhibitor used clinically to treat certain cancers, significantly reduces disease severity and parasite burden in *Leishmania*-infected mice, even if it is started significantly after lesions develop. Our results show that maximal *Leishmania* infection requires MAPK/ERK and highlight the potential for MAPK/ERK-mediated signaling pathways to be novel therapeutic targets for leishmaniasis.

**Lay summary:** *Leishmania* is a single-celled parasite that causes skin ulcers or a disseminated disease in humans. Our goal is to identify new drugs to treat *Leishmania* infection. *Leishmania* must live inside human immune cells to cause disease. If *Leishmania* is not able to enter human immune cells, it dies. Our studies demonstrate how *Leishmania* infection permits a set of proteins called MAP kinases to pass signals from one protein to the next within mammalian immune cells. The resulting signals allow *Leishmania* to enter into these immune cells and survive within its host. Importantly, trametinib, a drug that prevents these signals from MAP kinases, decreases the development of skin ulcers when it is given to mice that are infected with *Leishmania*. Our findings suggest that leishmaniasis could be treated with drugs that act on kinases found in humans rather than the parasites themselves.

**One sentence summary:** An Abl2-SYK-Raf-MEK-ERK pathway facilitates *Leishmania* uptake by phagocytic cells and promotes disease severity in *Leishmania*-infected mice.

## 1. INTRODUCTION

*Leishmania* spp. are obligate intracellular, protozoan parasites spread by the sand fly vector [1], causing leishmaniasis, a group of diseases ranging from self-healing cutaneous ulcers [2] to deadly disseminated visceral disease [3]. Leishmaniasis represents a major global health problem, since over 12 million people currently suffer from this disease. Approximately 2 million people are newly infected by *Leishmania* spp. annually [4]. Moreover, over 350 million people in 98 countries worldwide are at risk of developing leishmaniasis [4, 5], and the disease is now spreading into previously non-endemic areas, including the United States [6, 7]. *Leishmania*, particularly the species causing cutaneous disease, are understudied, and current treatments are few, expensive, poorly effective, and toxic. Resistance is also emerging [8–11]. Thus, identifying new drugs with novel mechanisms of action that circumvent resistance mechanisms is an urgent priority for effective drug development.

When an infected sand fly encounters its host, *Leishmania* promastigotes must be internalized by phagocytic cells such as macrophages (Mϕ) to persist in the host [12, 13]. This process is then repeated as the mammalian life cycle stage, the amastigote, replicates and spreads to other phagocytic cells to cause pathogenesis [14]. Amastigotes that do not enter phagocytic cells are thought to be cleared by the host immune system and/or die from lack of essential nutrients [14]. Major advancements have been made in understanding the immunomodulatory events involved in the pathogenesis of leishmaniasis. However, little is known about the basic molecular events underlying the pathogenesis, especially the host cellular signaling pathways, which facilitate the uptake of *Leishmania* by Mϕs and other phagocytes.

Based on prior studies, the Fcγ classes of receptors (FcγR) are the primary receptors on the host cell surface that permit amastigote uptake [14], and the complement receptor CR3 is the primary mediator of promastigote uptake [15]. Our previous studies have identified the Src [16] family kinases (SFK, and specifically Hck, Fgr, and Lyn), Abl1 (Abl) and Abl2 (Arg) [17] kinases, and Spleen Tyrosine Kinase (SYK) [18] as major protein families involved in phagocytosis and *Leishmania* uptake. In particular, we have demonstrated that Abl facilitates complement-mediated phagocytosis, and Arg enhances immunoglobulin/ FcγR-mediated phagocytosis [16, 17]. Modulation of these kinases via genetic manipulation or small molecule inhibitors abrogates *L. amazonensis* internalization by Mϕs, and thus limits disease progression in mice [16, 17]. However, our studies also indicate that these kinases have a limited role in the host’s immune response to *Leishmania* or parasite clearance [16, 17]. A drug that simultaneously decreases parasite uptake and promotes host-advantageous immunological responses could provide reductions in infection and disease burden. Host-cell active agents also could be used to combat other intracellular pathogens that employ similar uptake mechanisms.

Exactly how SFK, Abl family kinases, and SYK would regulate the events that occur during *Leishmania* uptake and phagocytosis has not been fully established. Others and we have demonstrated that CrkII is activated after signals have been relayed from SFK to Arg to SYK, leading to actin-rich phagocytic cup formation and pathogen uptake [16–19]. Interestingly, studies have been published primarily in cancer model systems indicating that SFK [20], Abl family kinases [20], and SYK [21, 22] also can activate MAPK/ERK signaling. MAPK/ERK are known to directly phosphorylate cytoplasmic or nuclear proteins, including transcription factors, that control several fundamental cellular processes from proliferation and differentiation to innate and adaptive immune responses [23–25]. In addition, Phorbol 12-myristate 13-acetate (PMA) is known to activate MAPK/ERK through protein kinase C (PKC) [26].

It is not clear whether MAPK/ERK signaling would mediate the uptake of *Leishmania* parasites by phagocytes. Research on *Plasmodium* indicated that inhibiting MEK1/2, which is upstream of ERK1/2, affects both adaptive and innate immune responses, which limits parasitemia and prevents pathogenesis [27]. Previous studies have also shown that MAPK/ERK signaling modulates the immune response to *L. amazonensis* infection to favor chronic infection, where subversion of MAPK/ERK signaling leads to parasite survival inside phagocytic cells, although these results vary depending on species [28–32]. Correspondingly, limiting MAPK activity using the older-generation MAPK inhibitors PD98059 or U0126 has been shown to enhance *L. amazonensis* killing by activating reactive oxygen species (ROS) and repressing IL-10 production [33–36]. These results implicate ERK1/2 phosphorylation as a mechanism through which *L. amazonensis* promotes persistence and subverts the immune response [33]. However, experiments characterizing the role of MAPK/ERK signaling during parasite uptake by Mϕ have not been conducted.

Here, we uncover an additional role for MAPK/ERK during *Leishmania* internalization by phagocytes. We show that Raf-1, MEK1/2, and ERK1/2 activity are necessary for efficient uptake of amastigotes. Host MAPK/ERK is activated by Arg and SYK to facilitate this process. In combination with prior results [16, 17, 33], our molecular studies elucidate an FcγR-SFK-Arg-SYK-Raf-MEK-ERK signaling relay as a major pathway involved in establishment of leishmaniasis by facilitating parasite uptake. In addition, using a mouse model, we establish that trametinib, a selective MEK inhibitor that is clinically approved to treat cancer, decreases footpad swelling and parasite burden in cutaneous leishmaniasis, even if it is started significantly after lesions are apparent. Our experiments highlight MAPK/ERK inhibitors as potential treatment modalities for this devastating disease.

## 2. MATERIALS AND METHODS

### 2.1 Mice

C57BL/6 mice were procured from Jackson Laboratory (Bar Harbor, ME). *Syk^flox/flox^ Cre^+^* and *Arg^−/−^Abl^flox/flox^ LysM Cre^+^* mice are either on a pure C57BL/6 background (SYK) [18] or have been backcrossed to C57BL/6 mice at least 5 times (Abl/Arg) [16], [17]. All the experiments involving knockout mice were conducted with WT littermates to ensure control for their genetic background. The Institutional Animal Care and Use Committee at University of Texas Southwestern Medical Center provided approval for all experimental protocols.

### 2.2 Mammalian cell culture

RAW 264.7 cells (ATCC, Manassas, VA) were cultured in Dulbecco’s modified Eagle’s medium (DMEM) plus 10% endotoxin-free, heat-inactivated, fetal bovine serum (FBS) (GeminiBio, Sacramento, CA) and 1% penicillin–streptomycin as described previously [16]. FBS was checked for LPS contamination using a highly sensitive Pierce™ Chromogenic Endotoxin Quant Kit (Thermo Scientific, Cat No. A39552), and was found to be endotoxin negative (ϕ 0.1 EU/ml). For experiments with bone marrow-derived Mϕs (BMDM), cells were obtained from the humeri, tibias, and femurs of *WT, Syk^flox/flox^ Cre^+^*, or *Arg^−/−^Abl^flox/flox^ LysM-Cre^+^* mice. They were differentiated to *WT, Syk^flox/flox^ Cre^+^*, or *Arg^−/−^Abl^flox/flox^ LysM-Cre^+^* BMDM by culturing them in DMEM, 10% FBS, and 20% L929 cell supernatant over 7 days. Differentiation to Mϕ was confirmed [17] and samples were screened for *Mycoplasma* contamination [37].

### 2.3 Parasite culture

*L. amazonensis* promastigotes (strain IFLA/BR/67/PH8, provided by Norma W. Andrews, University of Maryland, College Park, MD) were maintained at 26°C in Schneider’s Drosophila medium (Sigma, S9895, St. Louis, MO) plus 15% FBS and 1% penicillin–streptomycin [16, 17]. *L. amazonensis* amastigotes were grown axenically at 32°C in Schneider’s Drosophila medium at pH 5.5, plus 20% FBS, 1% penicillin–streptomycin, 0.1% hemin (25 mg/mL in 50% triethanolamine), 10 mM adenine, 5 mM L-glutamine, 0.25% glucose, 0.5% trypticase, and 40 mM sodium succinate (all from Sigma) [16, 17]. For uptake (internalization of amastigotes and promastigotes) assays, promastigotes were incubated for 7 days to maximize infective metacyclics (*i.e*., those isolated through a step Percoll gradient; Sigma). Amastigotes typically were grown axenically for 5 days prior to use [17]. To assess for amastigote growth and replication defects during incubation in MEK/ERK activators or inhibitors, medium containing 10 μM trametinib (Selleck Chemicals, Houston, TX), PMA (Selleck Chemicals), or 0.1% dimethyl sulfoxide (DMSO, Sigma) was used. To ensure infectivity, parasites were periodically passed in C57/BL6 mice prior to isolation of amastigotes from footpad lesions.

For some experiments, a transgenic stable mNeonGreen expressing line of *L. amazonensis* parasites (*L. amazonensis*-mNeon), containing a marker for hygromycin resistance, was generated [18]. *L. amazonensis*-mNeon promastigotes and amastigotes were maintained as above, but the media was supplemented with 100 μg/mL hygromycin (Sigma). *L. amazonensis* virulence was maintained by passage in C57BL/6 mice [17]. *L. amazonensis*-mNeonGreen parasites were detected with an excitation wavelength of 490–494 nm and emission of 515 nm.

### 2.4 *Leishmania* uptake assays

Experiments were adapted and modified as described previously [16, 17]. BMDM (*WT, Syk^flox/flox^ Cre^+^*, *Arg^−/−^Abl^flox/flox^ LysM-Cre^+^*) or RAW 264.7 cells were plated and treated with inhibitors (imatinib [Arg/Abl inhibitor], PD0325901 [MEK1/2 inhibitor], SCH772984 [ERK1/2 inhibitor], trametinib [MEK1/2 inhibitor]) (Selleck Chemicals, Houston, TX), PMA [ERK1/2 activator] or DMSO, where appropriate. Metacyclic promastigotes were incubated in fresh mouse serum for C3bi opsonization, and amastigotes were coated with anti-P8-proteoglycolipid complex (monoclonal antibody IgG1) [18]. Next, samples with RAW 264.7 cells or BMDMs were incubated with C3bi-opsonized promastigotes at 10 parasites to 1 Mϕ or with IgG-opsonized amastigotes at 10 parasites to 1 Mϕ for 2 hours (h), unless otherwise indicated, and fixed with 3% formaldehyde for 20 minutes (m). Mouse anti-gp46 antibody was allowed to bind to external promastigotes, whereas mouse anti-P8 antibody was incubated with external amastigotes, as described previously [16, 17]. Samples were then incubated with Alexa-Fluor-568-conjugated donkey anti-mouse-IgG secondary antibody. Any internalized parasite did not receive any antibody tagging and fluoresces green. Conversely, external parasites are visualized in both the red and green channels (orange in a merged image). All antibodies are listed in **Supplemental Table 3**. Thus, this three-color immunofluorescence assay [18] allowed distinguishing of internal (green) and external *L. amazonensis* promastigotes or amastigotes (green + red).

For quantification, samples were imaged with a BioTek Cytation 5 (Agilent, Santa Clara, CA) confocal imaging reader using a 40x objective lens with automated analysis performed [16]. At least 16 randomly selected fields were automatically visualized per well with each field assessing the number of Mϕs and amastigotes or promastigotes (internalized versus external). The absolute phagocytic index (the number of internalized amastigotes or promastigotes per 100 Mϕs) and adhesion index (the total number of amastigotes or promastigotes per 100 Mϕs) was calculated for each condition. The absolute phagocytic index or adhesion index for DMSO-treated conditions (control) was taken as the maximum value (100%) for each experiment, and the phagocytic index for imatinib, SCH772984, trametinib, or PMA-treated samples was determined and normalized relative to that of the control. Means ± SD were calculated. Each experiment was performed with at least 3 biological repeats and contained 3 technical replicates. A one-sample Student’s *t-* test or two-way ANOVA was used to determine statistical significance, where appropriate. The representative images shown were visualized using a Biotek Cytation 5 imaging multimode reader. Parasites in representative images were selected from files and linearly processed in Adobe Photoshop CS6 (version 13.0.6). Where applicable, permeabilized samples were labeled with far-red phalloidin (A22287, Thermo Fisher Scientific) at 1:100. For phagocytic cup analysis and representative images, data were collected on a Zeiss LSM 880 inverted confocal Airyscan microscope at 40X (analysis) or 63X (representative images). Images were created via Image J (1.52a, http://imagej.nih.gov/ij) and processed in Adobe Photoshop 13.0.6. [38].

### 2.5 Phagocytosis of beads

Experiments were adapted and modified as described previously [16, 17]. RAW 264.7 cells were plated at 70% confluence and incubated overnight in serum-free medium. Except where indicated, samples were pre-incubated in medium containing 3.3 μM imatinib, 1 μM SCH772984, 1 μM trametinib, 250 nM PMA or 0.1% DMSO for 1-2 h. Next, latex yellow–green 2 μm beads (Sigma) were coated with human IgM (Sigma, Cat. #: I-8260) and incubated in rabbit anti-IgM (Sigma, Cat. #: 270A) for IgG opsonization (confirmed as described previously [17]). Mϕs were incubated with 10–15 beads/ Mϕ for 2 h, unless otherwise stated, at 37°C. They were fixed with 3% formaldehyde for 20 m, blocked using 2% BSA in PBS (no permeabilization), and then incubated with rabbit anti-human IgM and Hoechst 33258 dye (Sigma) to visualize nuclei. Next, samples were incubated with an Alexa-Fluor-568-conjugated goat anti-rabbit-IgG secondary antibody (Invitrogen, Cat.#: A11034). All antibodies are listed in **Supplemental Table 3**. This multi-color immunofluorescence assay [39] allowed distinguishing of internalized (green) and external beads (green + red), similar to described above for internal vs external *L. amazonensis*. Quantification was done as above with a BioTek Cytation 5 confocal imaging reader using a 40x objective lens with automated analysis performed by the Cytation 5.

### 2.6 Western Blot

To analyze protein/phosphoprotein expression, western blotting was performed. Briefly, RAW 264.7 cells or BMDMs from *WT, Syk^flox/flox^ Cre^+^*, or *Arg^−/−^Abl^flox/flox^ LysM-Cre^+^* mice were incubated overnight in serum-free media (RAW 264.7 cells) or L929-free media (BMDM). Experiments were performed at ∼70% confluence. Where appropriate, Mϕ were pre-incubated in imatinib, entospletinib, PD0325901, selumetinib, SCH772984, trametinib, PMA or DMSO. IgG-opsonized amastigotes or IgG-opsonized beads were added to adherent starved Mϕs for 30 m, unless otherwise indicated [40], and cells lysed with RIPA extraction buffer and halt protease and phosphatase inhibitor (Thermo Scientific, Cat. #: 78440, Richardson, TX). Protein purity and concentration were addressed using Coomassie Blue staining and BCA assay, respectively, following the manufacturers’ protocols. Total proteins were resolved by polyacrylamide gel electrophoresis (12% gels) and transferred to PVDF membranes (BioRad). The membranes were blocked in 5% non-fat dry milk in Tris-buffered saline (TBS) (20 mM Tris at pH 7.6, 150 mM NaCl). The membrane was incubated overnight at 4°C with the primary antibody in 5% BSA and TBS-T. Next, horseradish peroxidase (HRP)-conjugated anti-mouse secondary antibody (#7076, Cell Signaling Technology) was added at 1:5000 in 5% non-fat dry milk in TBS-T for 1 h. All antibodies are provided in **Supplemental Table 3**. Membranes were incubated in SuperSignal West Pico Chemiluminescent Substrate (Thermo Scientific) for 5 m and visualized by using a phosphorimager (ImageQuant LAS 4000, GE Healthcare). For analysis, relative amounts of phosphorylated proteins were compared to total proteins with ImageJ analysis software (https://imagej.nih.gov/ij); membranes were stripped and reprobed). Each experiment had at least 3 biological replicates. Two-way ANOVA was used for statistical tests.

### 2.7 Leishmania animal model

For the mouse footpad model of cutaneous leishmaniasis, 10 female C57BL/6 mice per group were infected between 6 and 8 weeks (w) of age as previously described [16, 17]. Briefly, 1×10^6^ metacyclic promastigotes were suspended in PBS and injected subcutaneously in the dorsal side of the right hind foot. For the experiments in Fig 6A-C, mice were provided 2 mg/kg body weight of trametinib or drug vehicle (4% DMSO and 1% corn oil) by oral gavage daily starting a w before infection and continuing until the mice were euthanized. Two independent experiments were performed. For the experiments in Fig 6D-F, mice were provided 2 mg/kg body weight of trametinib or drug vehicle by oral gavage daily starting 6 w after infection and continuing until the mice were euthanized. Two independent experiments were performed. Lesion growth was documented with calipers every other w by an investigator who was blinded to condition. The ratio of the infected: uninfected foot size was obtained [41]. ANOVA was used to compare diluent and trametinib-treated mouse groups to swelling at time 0. The number of parasites in lesions was determined by limiting dilution upon termination of the experiment between w 14 to 16 [42].

Lymph nodes from infected diluent versus trametinib-treated mice were obtained for cytokine and chemokine profiling [42, 43]. Briefly, lymph node cells were incubated in RPMI 1640 (Invitrogen) plus 10% heat-inactivated FBS, 5 × 10^−5^ M 2-mercaptoethanol (Sigma), and 1% penicillin-streptomycin. Cells were stimulated with promastigote lysates (equivalent to 1 × 10^6^ or 1 × 10^5^ parasites per sample, as indicated) or concanavalin A (ConA) (5 μg/ml; Sigma). Supernatants were harvested at 72 h and the levels of the cytokines and chemokines IL-4, IL-10, IL-12, IL-13, IL-17, IFN-γ, IL-1β, IL-1α, IP-10, MIP-1α (CCL3), MIP-2 (CXCL2), CXCL9, RANTES (CCL5), and MIP-1β (CCL4) were profiled using the custom multiplex Luminex Platform (R&D Systems, Inc, Minneapolis, MN) through the UT Southwestern Microarray Core as described previously [44, 45]. Background cytokine levels were calculated using unstimulated supernatants.

## 3. RESULTS

### 3.1 MAPK/ERK signaling in Mϕ facilitates *Leishmania* uptake

To directly test whether the MAPK/ERK pathway mediates *Leishmania* uptake, we first explored the effect of the MEK1/2 inhibitor trametinib on the uptake of *L. amazonensis*. We can grow both promastigotes and amastigotes of this species in tissue culture without host cells (*i.e*., axenic culture), which facilitates our studies of *Leishmania* uptake by Mϕs. Trametinib is a highly selective and potent MEK1/2 inhibitor, with an IC_50_ for the purified kinase ranging from 0.92 nM to 3.4 nM in cell-free assays [46, 47]. Trametinib has no effect on the kinase activity of c-Raf, B-Raf, or ERK1/2. We pretreated RAW 264.7 cells, a Mϕ−like cell line, for 2 hours (h) with trametinib or imatinib, an Abl/Arg inhibitor with an IC_50_ of 100-500 nM in kinase assays [35], as a positive control. We then added *Leishmania* to samples to allow uptake and processed for immunofluorescence microscopy. Visually, trametinib decreased the uptake of IgG-opsonized amastigotes in RAW 264.7 cells, comparable to the reductions seen after treatment with imatinib (**Fig. 1A**). Defects in internalization were quantified by calculating the phagocytic index (PI), or number of internalized parasites per 100 Mϕ, which was lower in trametinib-treated samples (**Fig. 1B**). No differences were seen in the total (external + internal) number of *Leishmania* per 100 RAW 264.7 cells, which is termed an adhesion index, among these conditions (**Fig. 1C**). We performed a titration of trametinib’s effects on amastigote uptake, and the maximal reduction in PI was shown at 1 μM trametinib (**Fig. 1D, E**). We also determined that over 72 h, the CC_50_ (cytotoxic concentration required to kill 50% of viable cells) for RAW 264.7 cells was greater than our maximum concentration of 2.5 μM. Internalization defects remained even after prolonged incubation in trametinib (up to 240 minutes, m) and were not due to reduced adhesion, as amastigotes continued to be bound to trametinib-treated RAW 264.7 cells at levels indistinguishable from controls (**Fig. 1F, G**). Treating Mϕs with other MAPK/ERK pathway inhibitors [48] such as selumetinib and PD0325901 (MEK1/2 inhibitors), as well as SCH772984 (an ERK1/2 inhibitor), also significantly reduced amastigote uptake without affecting the total number of bound parasites (**Supplemental Fig. 1A, B**). In addition, we found that treating RAW 264.7 cells with the MEK1/2 inhibitor PD0325901 [49] decreased the internalization of promastigotes to a degree that was comparable to imatinib (**Supplemental Fig. 1C, D**). Next, we characterized growth curves and found that trametinib showed no impact on cell viability and motility of parasites in culture (**Fig. 1H**). Together, these similar results using multiple MAPK/ERK inhibitors establish that targeting MEK1/2 and ERK1/2 significantly attenuates internalization of both *L. amazonensis* life cycle stages. In addition, as previously published for MAPK/ERK inhibitors, amastigotes within Mϕs had a ∼50% reduction in survival when cultures were treated with trametinib for 72 h (**Supplemental Fig. 1E**).

**Figure 1.**
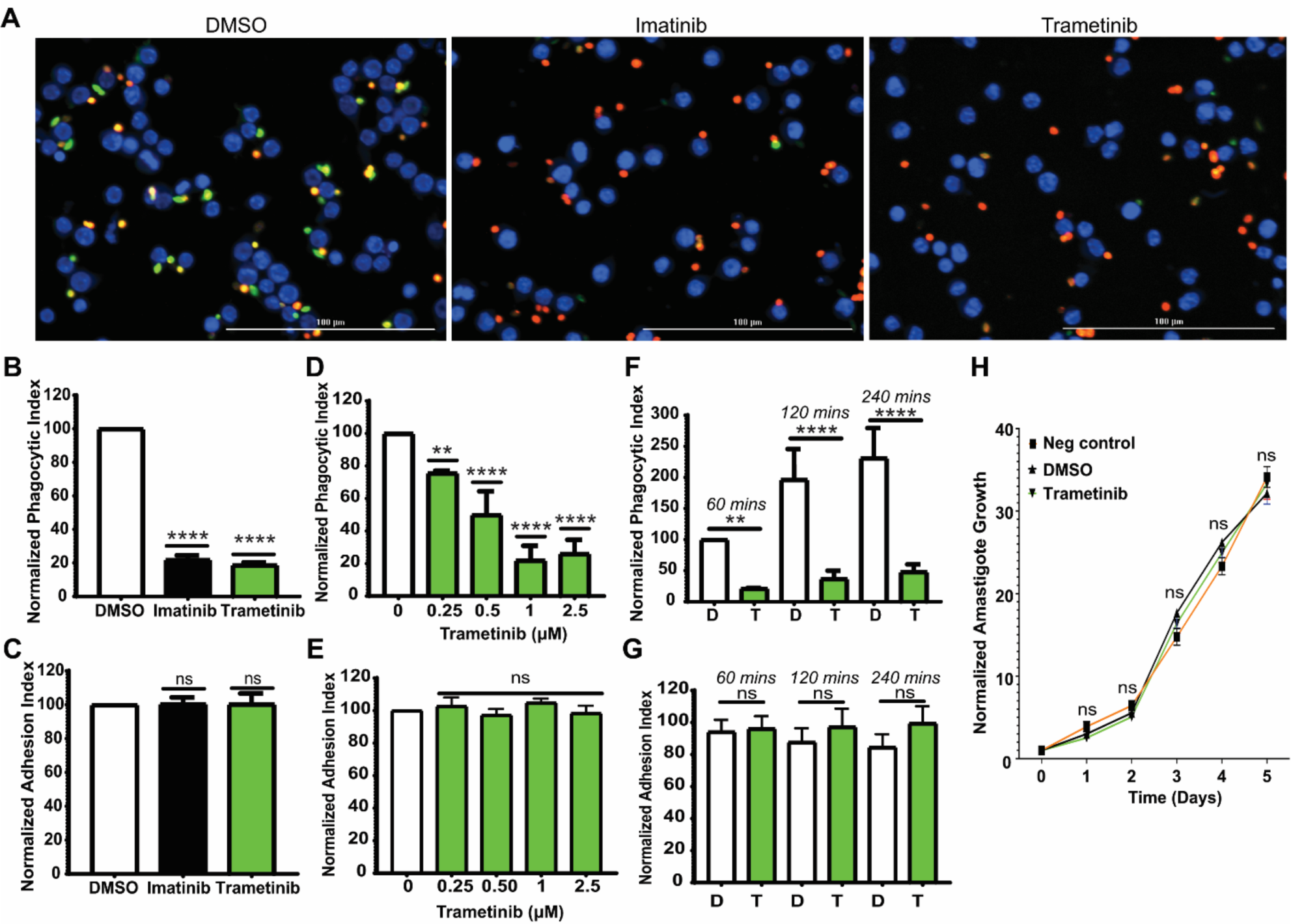
The MAPK/ERK pathway is required for optimal amastigote uptake. **(A)** Trametinib decreases IgG-opsonized amastigote uptake. RAW 264.7 cells were treated with 3.3 μM imatinib, 1 μM trametinib or DMSO for 2 h, and 10 IgG-opsonized amastigotes were incubated per RAW 264.7 cell for 2 h at 37°C. An immunofluorescence assay distinguished between intracellular (green) and extracellular (red + green) amastigotes. Nuclei are labeled with Hoescht 33342 (blue). Scale bar, 100 μm. **(B)** Graph showing the mean phagocytic index (PI) ± standard deviation (SD) for RAW 264.7 cells normalized to DMSO for each experiment (100%), quantified from n = 6 biological experiments. Comparisons between imatinib and trametinib by ANOVA were nonsignificant. **(C)** Imatinib and trametinib do not affect the total (external + internal) number of amastigotes per 100 RAW cells (termed an adhesion index). Shown are percentages of total amastigotes per 100 imatinib-treated and trametinib-treated RAW 264.7 cells relative to DMSO-treated cells for the experiment shown in B. **(D)** Titration of trametinib demonstrates that the reduction of amastigote internalization is maximal at ∼100x the purified MEK1/2 IC_50_. RAW 264.7 cells were treated with increasing doses of trametinib (0.25, 0.5, 1, or 2.5 μM) or DMSO. Bar graph shows the mean PI ± SD for trametinib-treated samples relative to DMSO-treated samples for each experiment (100%) for 4 biological experiments. **(E)** The adhesion index for amastigotes from experiment in D is not affected by increasing doses of trametinib. Experiment performed as in C. **(F)** The relative decrease in PI for trametinib-treated samples persists even after prolonged incubation. RAW 264.7 cells were treated with DMSO (D) or 1 μM trametinib (T) prior to addition of IgG-opsonized amastigotes for 1 h, 2 h, or 4 h, as indicated. Shown is the mean PI ± SD for each time point for the trametinib-treated condition, relative to the DMSO-treated condition, calculated from 4 biological experiments. **(G)** Treating samples with trametinib for prolonged periods does not affect the adhesion index for amastigotes. Shown are the total number of amastigotes (internal + external) per 100 RAW 264.7 cells for the experiments performed in F. For all experiments, *\*\* P* < 0.01, **** *P* < 0.0001, ns = nonsignificant compared to the relevant DMSO-treated category by ANOVA. **(H)** Trametinib is not toxic to amastigotes. Shown is one representative experiment of five experiments with two technical replicates following the number of amastigotes growing in untreated, DMSO-treated, or trametinib-treated media over 5 days relative to on day 0. ns, *P* > 0.05 compared to DMSO category by ANOVA.

### 3.2 cRaf, MEK1/2, and ERK1/2 are activated during amastigote uptake

Having established that MAPK/ERK are important for *Leishmania* uptake, we assessed whether kinases in this pathway were activated in host cells during amastigote internalization. Raf-1 (cRaf), MEK1/2, and ERK1/2 are key members of the MAPK/ERK pathway [25, 50]. We thus measured cRaf, MEK1/2, and ERK1/2 phosphorylation in RAW 264.7 cells to examine the relative phosphorylation level of each kinase during internalization of amastigotes. Essentially no phosphorylated ERK or ERK was detected in amastigote samples (**Supplemental Fig. 2A**). However, amastigotes potently induced ERK1/2 phosphorylation in Mϕs, with phosphorylation levels increasing by 3 and 4.5-fold at 10 and 30 m post-infection, respectively (**Fig. 2A**). Similarly, MEK1/2 (**Fig. 2B**) and cRaf (**Fig. 2C**) were activated by amastigotes that were entering host cells. These data indicate that amastigotes rapidly activate MAPK/ERK in a sustained fashion during their uptake by Mϕs.

**Figure 2.**
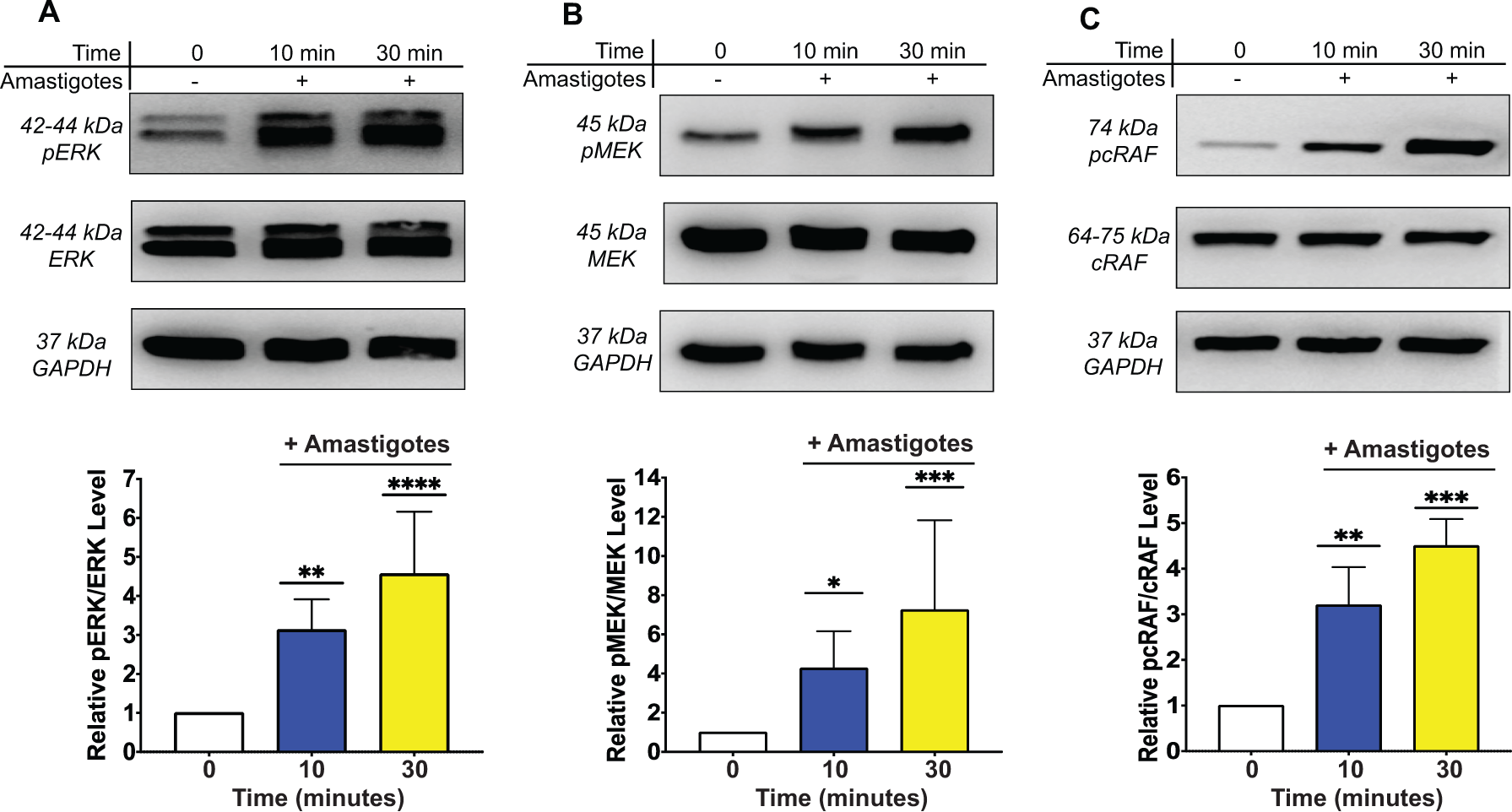
MAPK/ERK are activated during *Leishmania* uptake by Mϕ. Amastigotes were added to ≥ 70% confluent RAW 264.7 cells. Equal amounts of protein from whole lysates were employed for each category. **Top:** Representative blots displaying relative activation of MAPK/ERK. Phosphorylated **(A)** ERK1/2, **(B)** MEK1/2, **(C)** cRaf (Raf-1) and the corresponding total proteins were detected by Western Blot. An unrelated protein, GAPDH, is shown as an additional loading control. **Bottom:** Quantification of phosphorylated MAPK/ERK levels. Graphs shown in **(A)** for ERK1/2, **(B)** for MEK1/2, and **(C)** for cRaf represent mean ± SD of relative phosphorylated kinase levels, normalized to total kinase levels, for at least 3 biological replicates per condition. * *P* < 0.05, ** *P* < 0.01, *\*\*\* P* < 0.001, **** *P* < 0.0001 by ANOVA compared to the 0 m time point for each set of kinases.

### 3.3 PMA rescues MAPK/ERK related uptake defects

PMA is known to activate ERK1/2 through PKC, with an IC_50_ of 11.7 nM against purified kinase [26]. Consistent with this, although PKC activation was seen during the uptake of amastigotes, PMA treatment increased PKC phosphorylation in RAW 264.7 cells to 8-fold higher levels (**Fig. 3A, B**). Importantly, 10 μM PMA did not affect growth of amastigotes in culture (**Fig. 3C**), indicating that it does not act directly on *Leishmania*. We next investigated whether PMA affected amastigote internalization. To perform these experiments, and to confirm our prior results from a cell line using primary cells, we isolated bone marrow derived-Mϕs (BMDM) from C57BL/6 mice. PMA-treated BMDM were able to internalize IgG-coated amastigotes at higher levels than DMSO-treated BMDM in a dose-dependent manner, reaching maximal internalization at 2.5 μM PMA (**Fig. 3D-F**). Furthermore, PMA treatment rescued trametinib inhibition of *Leishmania* uptake in BMDM to PI that were indistinguishable from those of DMSO-treated controls (**Fig. 3G, H**). Collectively, our data indicate that ERK1/2 kinase activity is necessary for maximal *Leishmania* uptake.

**Figure 3.**
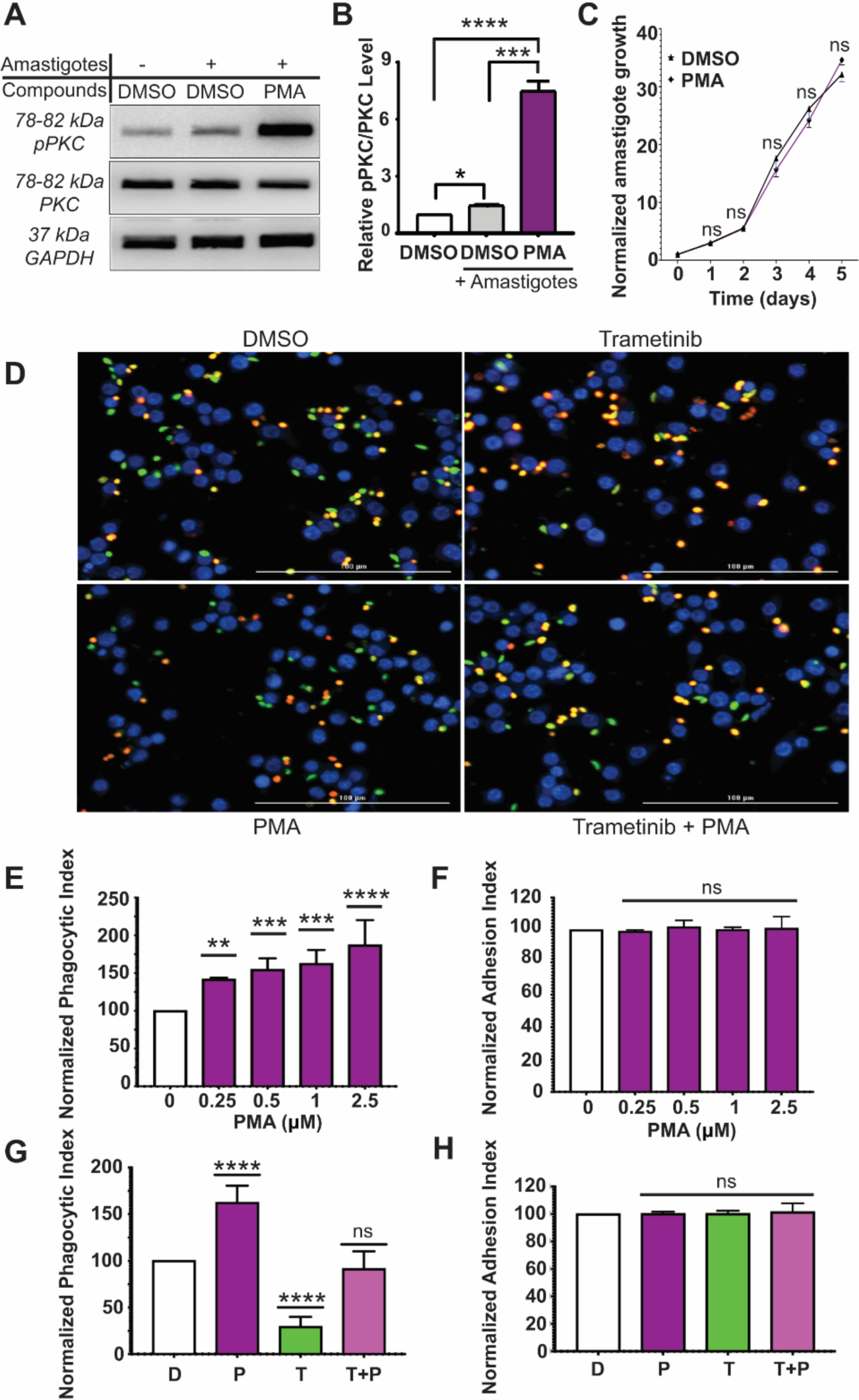
PMA activates ERK1/2 to modulate amastigote uptake. **(A, B)** PKC typically is only slightly activated during amastigote uptake. PMA serves as a positive control. Amastigotes were added to monolayers of RAW 264.7 cells and processed as above. Shown are **(A)** representative blots demonstrating and **(B)** bar graphs quantifying relative activation of pPKC/PKC. *\* P* < 0.05*, *** P* < 0.001, *\*\*\*\* P* < 0.0001 compared with samples indicated by brackets (ANOVA). n = 3 experiments. **(C)** PMA is not toxic to amastigotes. Shown is one representative experiment of five experiments with two technical repeats following the number of amastigotes growing in DMSO-treated or PMA-treated media over 5 days relative to on day 0. ns, *P* > 0.05 compared to DMSO category by ANOVA. **(D)** Representative image demonstrating that PMA rescues trametinib inhibition of *Leishmania* amastigote uptake. BMDM were treated with DMSO, 1 μM PMA, 1 μM trametinib or a combination of 1 μM trametinib and 1 μM PMA prior to incubation with IgG-opsonized amastigotes for 2 hours. **(E)** PMA increases IgG-opsonized amastigote uptake. BMDM were treated with DMSO or increasing doses of PMA (0.25, 0.5, 1, 2.5 μM) prior to incubation with IgG-opsonized amastigotes for the times indicated. n = 4 experiments. *\*\* P* < 0.01*, *** P* < 0.001, **** *P* < 0.0001 compared to DMSO category by ANOVA. **(F)** PMA does not affect the adhesion index for amastigotes. ns, not significant by ANOVA. **(G)** Mean PI ± SD for BMDM treated with trametinib/PMA relative to DMSO-treated samples for each experiment (100%), obtained from 3 biological experiments. **** *P* < 0.0001, ns = nonsignificant by ANOVA compared to DMSO-treated sample. **(H)** Trametinib/PMA does not affect amastigote adhesion for the experiment shown in G.

### 3.4 ERK1/2 is activated by Arg/Abl and SYK during amastigote uptake

Next, we investigated the mechanism by which *L. amazonensis* amastigotes induced ERK1/2 phosphorylation during their uptake. Previous research has shown that Arg facilitates *Leishmania* amastigote uptake downstream of SFK [16]. In addition, inhibition of SYK, a tyrosine kinase implicated in FcγR immunoglobulin-mediated signaling [18, 33, 51, 52] reduces ERK1/2 activation in a dose-dependent manner in other systems [18, 33]. To test for roles for these kinases, small molecule inhibitors (imatinib, entospletinib [IC_50_ for activity against purified SYK = 7.7 nM] [18], SCH772984, trametinib), and PMA were added to RAW 264.7 cells before infection with amastigotes. Inhibition of Arg and SYK with imatinib and entospletinib, respectively, significantly reduced ERK1/2 phosphorylation, to a similar degree to the MAPK/ERK inhibitors SCH772984 and trametinib (**Fig. 4A, B**). As expected, PMA, the positive control, significantly increased ERK1/2 phosphorylation (**Fig. 4A, B**). The inhibitors and activators followed similar trends for promastigote-induced ERK activity (**Supplemental Fig. 2B**). Thus, we could modulate *Leishmania*-induced ERK activity by both inhibitors and activators *in vitro*.

**Figure 4.**
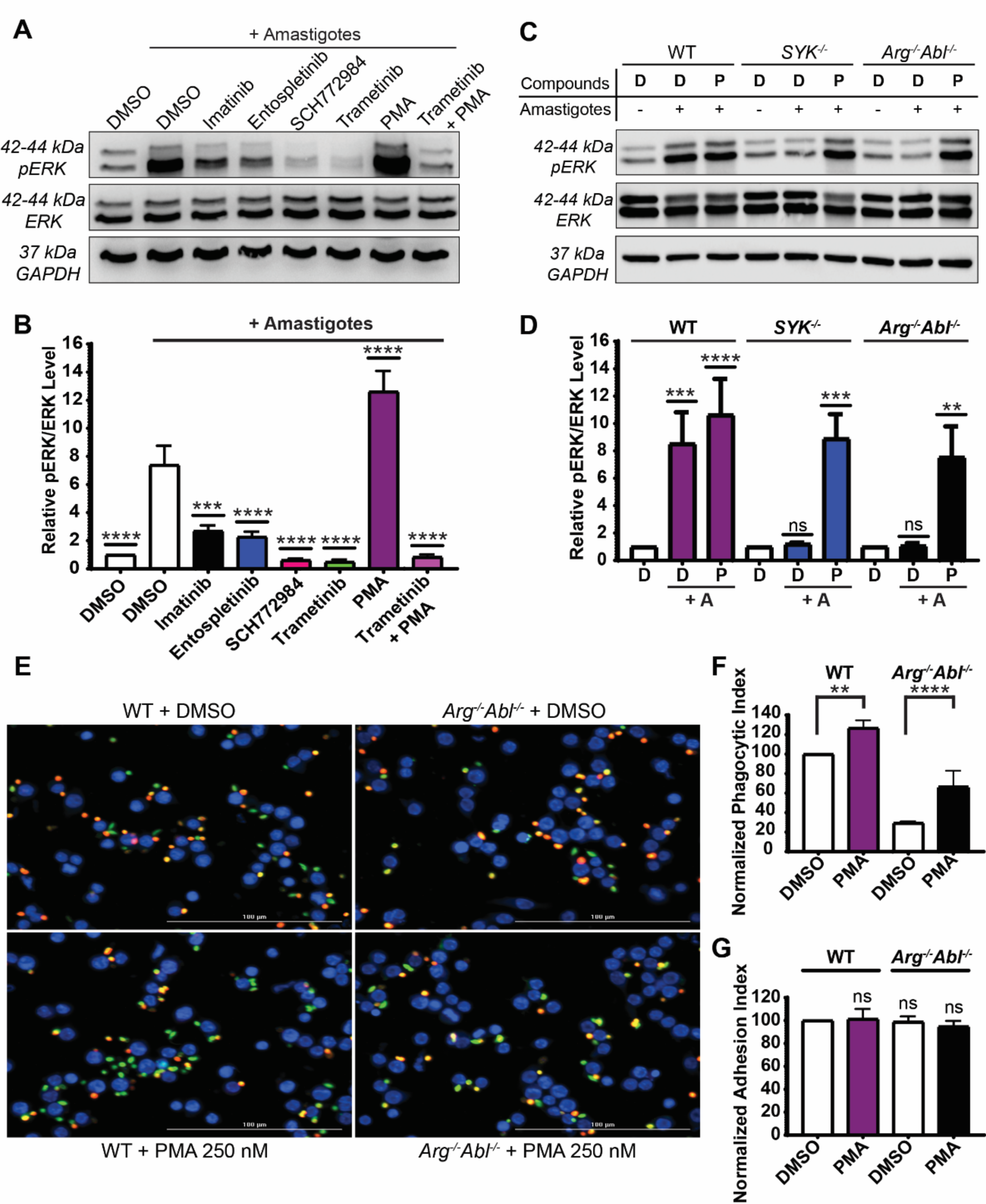
SYK and Arg/Abl stimulate ERK1/2 phosphorylation during amastigote uptake. **(A)** Phosphorylation of ERK1/2 in RAW 264.7 cells is induced upon amastigote uptake and is decreased by imatinib (Arg/Abl inhibitor), entospletinib (SYK inhibitor), SCH772984 (ERK1/2 inhibitor), and trametinib (MEK1/2 inhibitor). PMA serves as a positive control for pERK stimulation. RAW 264.7 cells were allowed to adhere to plates, and then treated with compounds or DMSO prior to IgG-opsonized amastigote addition, lysis, and immunoblotting. Representative immunoblots of pERK1/2 (top) and total ERK1/2 (bottom) [with (+) or without (–) amastigotes] are shown. **(B)** Relative pERK levels, normalized to ERK1/2 levels, among categories shown in A. Levels of pERK are normalized to amastigote-exposed DMSO-treated RAW 264.7 cells (100%). 5 biological replicates are quantified. *\*\*\* P* < 0.001, *\*\*\*\* P* < 0.0001 compared with amastigote-stimulated, DMSO-treated RAW 264.7 cells (ANOVA). **(C)** Phosphorylation of ERK1/2 induced upon FcγR engagement is decreased in *SYK*^−/−^ and *Arg*^−/−^*Abl*^−/−^ BMDM compared to WT BMDM. Shown is a representative immunoblot of pERK1/2 (top) and total ERK1/2 (bottom) in WT, *SYK*^−/−^ and *Arg*^−/−^*Abl*^−/−^ BMDM treated with DMSO (D) or PMA (P), ± incubation with amastigotes. **(D)** Graph showing relative levels of pERK1/2, normalized to ERK1/2 levels, among WT, *SYK*^−/−^, and *Arg*^−/−^*Abl*^−/−^ BMDM treated with DMSO or PMA (P), ± amastigotes (A). *\*\* P* < 0.01, *\*\*\* P* < 0.001, *\*\*\*\* P* < 0.0001, ns: not significant by ANOVA (n = 6 experiments), compared to DMSO + amastigote category for each condition. **(E-G)** Primary cells lacking Arg and Abl exhibit decreases in amastigote uptake, but PMA can partially rescue this amastigote internalization defect. **(E)** Representative image performed as in Fig 1. **(F)** Mean phagocytic index ± SD for PMA-treated and/or *Arg*^−/−^ *Abl*^−/−^ BMDM relative to WT DMSO-treated BMDM. **(G)** DMSO and PMA do not affect the adhesion indexes for the experiments shown in F. n for F-G = 3 experiments. *\*\* P* < 0.01; *\*\*\* P* < 0.001; *\*\*\*\* P* < 0.0001; ns, not significant compared with DMSO-treated WT BMDM (ANOVA).

To further characterize the role of SYK and Arg in *Leishmania*-induced ERK phosphorylation, we next isolated BMDM from mice lacking Abl family kinases (*Arg^−/−^Abl^flox/flox^LysM Cre^+^* [hereafter termed *Arg^−/−^Abl^−/−^* when discussing isolated BMDM]), SYK (*SYK^flox/flox^ Cre^+^*, abbreviated *SYK^−/−^* when referring to isolated BMDM) or wild type (*WT*) littermates. We have previously shown that amastigote uptake is dependent upon Arg but not Abl [15–16]. Our immunoblots showed *Arg^−/−^Abl^−/−^*and *SYK^−/−^* BMDM nearly completely lack Arg or SYK, respectively, while ERK1/2 expression was maintained (**Supplemental Fig. 2C**). Consistent with our chemical inhibition experiments, *SYK*^−/−^ and *Arg^−/−^Abl^−/−^* BMDM limited amastigote-mediated ERK1/2 phosphorylation, demonstrating that SYK and Arg/Abl are necessary for stimulating ERK1/2 phosphorylation following engagement of the FcγR (**Fig. 4C, D**). At low doses (250 nM), PMA rescued ERK1/2 activity in *Arg^−/−^Abl^−/−^* and *SYK^−/−^* BMDM. This suggested 1) amastigotes operate through a distinct pathway to activate ERK1/2 compared to PMA, which activates MAPK/ERK through PKC [26] and 2) during FcγR-mediated amastigote infection, Arg and SYK family kinases are upstream of MAPK/ERK and are necessary to activate the MAPK/ERK pathway.

To directly test whether Abl family kinases are critical for ERK1/2-mediated amastigote uptake, we treated *WT* and *Arg^−/−^Abl^−/−^* BMDM with DMSO or PMA prior to infection with IgG-opsonized amastigotes. We found that *Arg^−/−^ Abl^−/−^* BMDM displayed defects in IgG-mediated amastigote uptake, while PMA treatment partially rescued these defects (**Fig. 4E-G**), returning uptake to 66.4 ± 16.6% (mean ± SD) compared to 29.3 ± 0.9% with DMSO. Similarly, SYK was necessary for efficient amastigote uptake, and the decreased PI of *SYK^−/−^* BMDM was partially rescued by PMA treatment (**Supplemental Fig. 2D, E)**. Taken together, these results, in combination with prior data, are consistent with a signaling pathway in which Arg/Abl and SYK act upstream of and activate the ERK1/2 pathway (which is at least partly independent from PKC) to facilitate amastigote uptake [26].

### 3.5 MAPK/ERK inhibition minimally impacts phagocytosis of opsonized beads

*Arg^−/−^* BMDM have been shown to have impaired FcR-mediated phagocytosis overall, as demonstrated using IgG-coated (opsonized) beads [17]. Thus, we hypothesized that the Arg-SYK-ERK pathway may be necessary for the general process of phagocytosis, in addition to amastigote uptake. To delineate the specific contributions of MAPK/ERK to phagocytosis, we examined bead uptake compared to amastigote uptake by RAW 264.7 cells. Surprisingly, phagocytosis of IgG-opsonized beads by RAW 264.7 cells was minimally affected by trametinib (**Fig. 5A, B**) or SCH772984 (**Supplemental Fig. 3),** both of which resulted in only a ∼30% reduction in uptake. Conversely, the uptake of IgG-coated amastigotes (**Fig. 1**) was reduced by ∼70% with trametinib treatment. However, when RAW 264.7 cells were treated with imatinib, similar decreases in phagocytosis of IgG-opsonized beads versus amastigotes were observed. These results suggest that Arg/Abl play a more generalized role in phagocytosis, as we have demonstrated previously [17]. Collectively, our data suggest that ERK1/2 activation is important for *Leishmania* uptake specifically but may play a minor role in standard phagocytic mechanisms.

**Figure 5.**
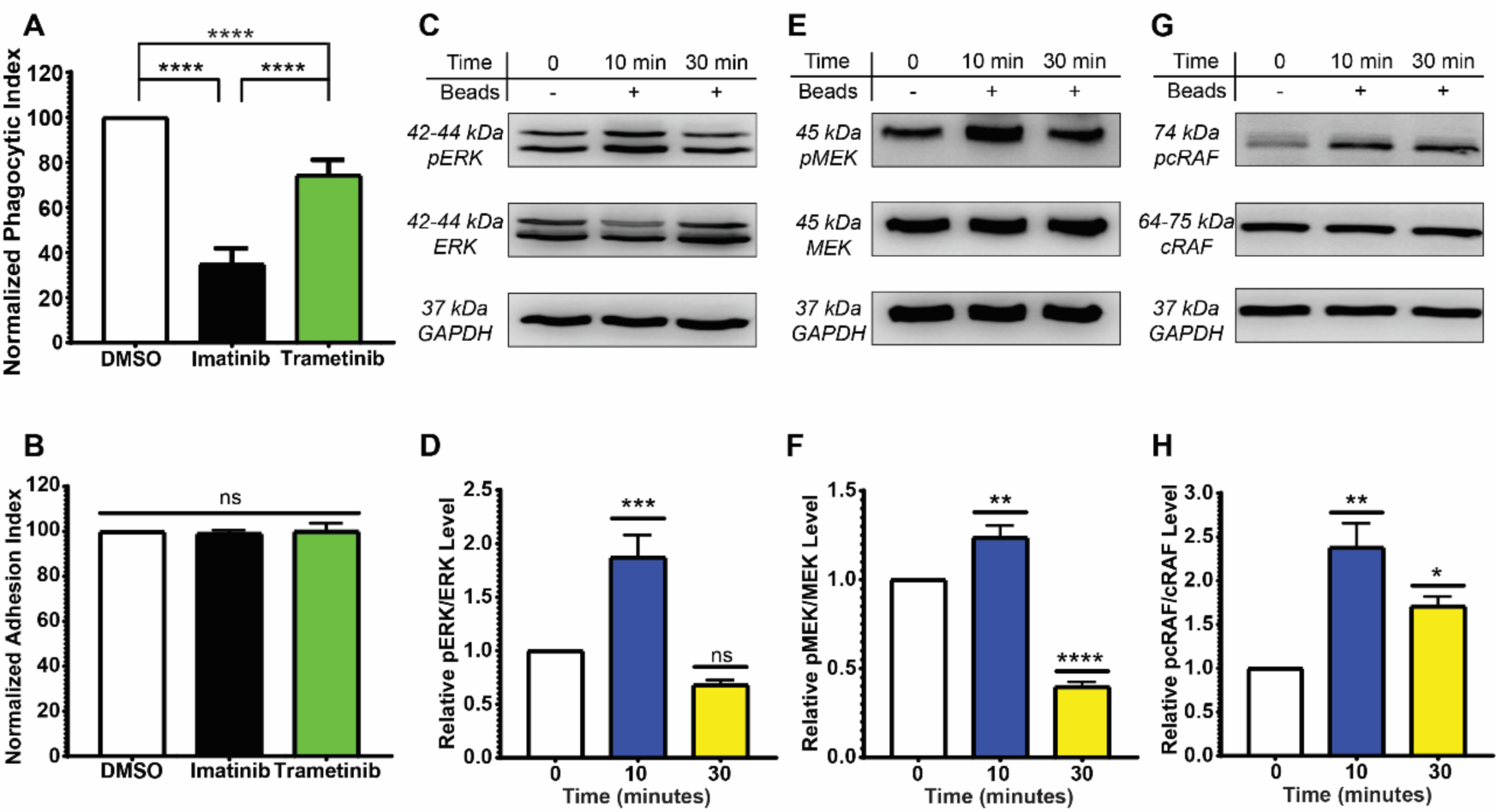
ERK 1/2’s role in *Leishmania* uptake is greater than for phagocytosis of beads. **(A)** Trametinib mildly decreases bead internalization. RAW 264.7 cells were treated with 3.3 μM imatinib, 1 μM trametinib or DMSO for 2 hr. prior to addition of 10 IgG-opsonized beads per RAW 264.7 cell for 2 h at 37°C. Treating RAW 264.7 cells with trametinib decreases uptake of IgG-opsonized beads to a lesser degree than imatinib. Graph shows the mean PI ± SD for each category, normalized to DMSO (100%). **** *P* < 0.0001, ns: not significant compared to indicated categories by ANOVA (n = 6 biological experiments). **(B)** Imatinib and trametinib does not affect the adhesion index for beads in the experiments shown in A. **(C-H)** IgG-coated beads activate the MAPK/ERK pathway, but phosphorylation is not sustained. Beads were added to > 70% confluent RAW 264.7 cells and experiments were processed as in Fig 2. Top: Phosphorylated forms of **(C)** ERK, **(E)** MEK1/2, **(G)** cRaf and the corresponding total kinase were detected by Western blotting. Bottom: Relative activation was quantified for **(D)** ERK, **(F)** MEK1/2, and **(H)** cRaf, using the methods described in Fig 2. * *P* < 0.05, ** P < 0.01, *\*\*\* P* < 0.001, **** *P* < 0.0001 and ns: not significant compared to 0 m timepoint by ANOVA (n ≥ 3 experiments).

**Figure 6.**
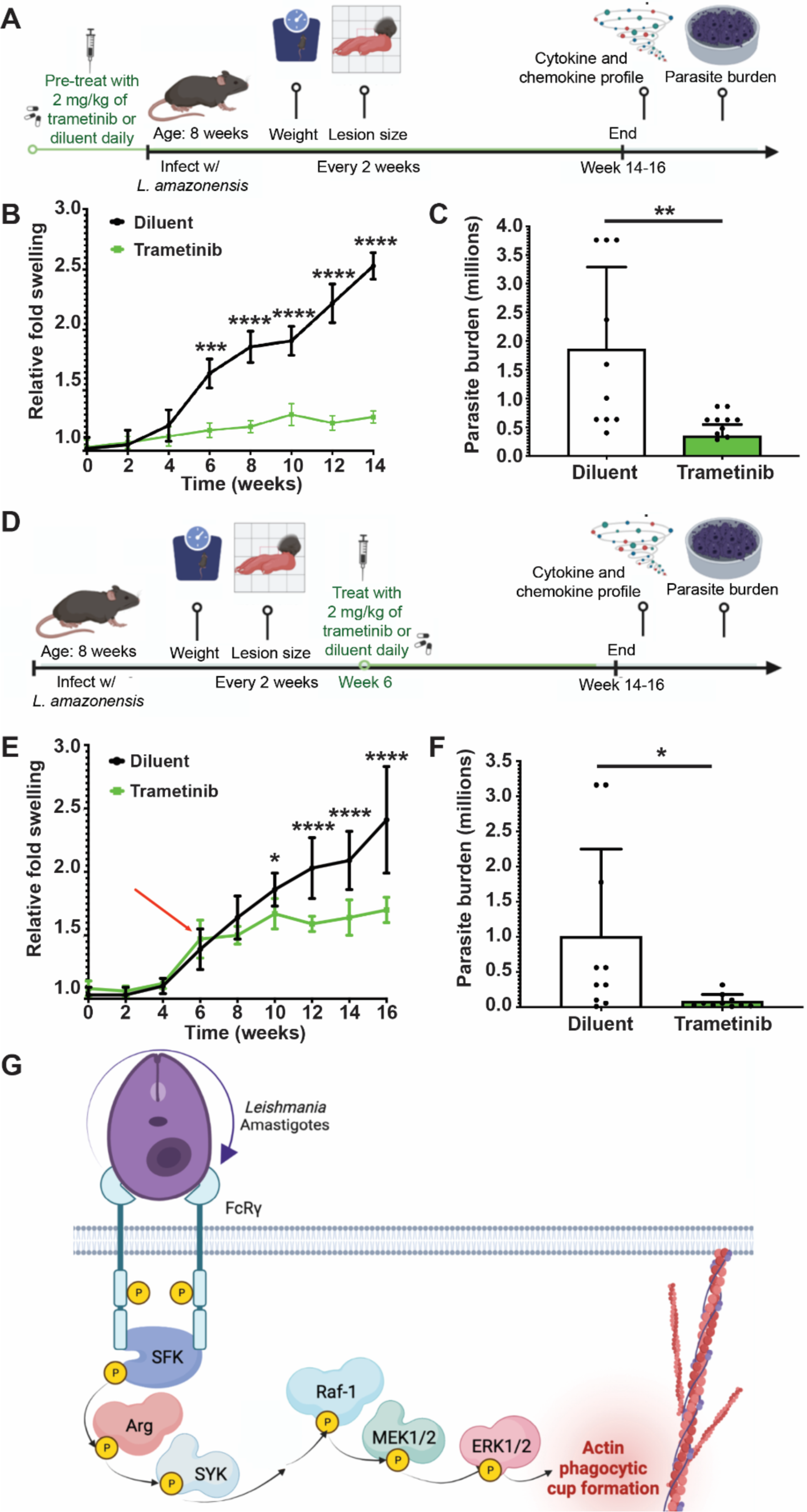
MAPK/ERK permit efficient infection in a mouse model of cutaneous leishmaniasis. **(A)** Mouse footpad model for cutaneous leishmaniasis. C57BL/6 mice diluent and trametinib-treated groups were injected with promastigotes in the hind footpad. Mice were given 2 mg/kg body weight of trametinib or an equal volume of diluent by oral gavage daily, starting 7 days before infection and continuing until the mice were euthanized. Two experiments were performed. Shown in the following panels is one representative experiment containing 10 female mice per group. **(B)** Trametinib given prior to infection reduces lesion size in mice. Depth of foot swelling was monitored with calipers by a blinded investigator. Results represent the mean ± SD foot size increase over the uninfected foot (normalized to 1). The control (diluent) and drug-treated groups were compared to swelling at time 0 by ANOVA; *\* P* < 0.05, *\*\*\*\* P* < 0.0001, ns, not significant. **(C)** Trametinib given prior to infection significantly reduces parasite burden. Parasite burdens in infected footpads were determined by limiting dilution as described in *Materials and Methods*. Plotted is the mean number of parasites (*i.e.*, parasite burden) in millions at the end of the experiment shown in B ± SD. In this example, there were 1.41 × 10^6^ parasites in the control mice versus 2.03 × 10^5^ parasites in the trametinib-treated mice. *\*P* < 0.05 (two-tailed *t*-test). **(D)** C57BL/6 mice diluent and trametinib-treated groups were injected with *L. amazonensis* promastigotes as above. Starting 6 weeks after inoculation (arrow), mice were given 2 mg/kg body weight of trametinib or vehicle by oral gavage daily and continuing until the experiment was terminated. (D) Trametinib given once lesions have begun reduces lesion size in trametinib-treated mice compared to diluent-treated mice. Arrow shows start time of drug treatment. Shown is one representative experiment containing 10 female mice per group. Statistics calculated as in B. **(F)** Lesions in mice treated with trametinib 6 weeks after infection contain fewer *Leishmania* parasites than control mice. Plotted is the mean ± SD lesion parasite burden in millions at the end of the experiment shown in E; in this example, there were 1.01 × 10^6^ parasites in the control mice versus 8.29 × 10^4^ parasites in the trametinib-treated mice (>10-fold reduction). \**P* <0.05 (two-tailed *t*-test). (**G**) MAPK/ERK signal transduction in Mϕ facilitates *Leishmania* uptake and pathogenesis. Upon FcγR ligation by amastigotes, SFK, Arg, and SYK are activated and relay signals to the MAPK/ERK pathway, which allows the internalization of *Leishmania*. Diagrams created with Biorender.

We also tested whether MAPK/ERK activity was induced following addition of beads compared to amastigotes. Beads induced ERK1/2 phosphorylation, but to levels that were lower than in seen during amastigote uptake (**Fig. 5C, D**). This phosphorylation occurred in a time-dependent fashion, such that activation reached maximal levels between 5 and 10 m, and ERK1/2 phosphorylation was not detectable after 30 m (**Fig. 5C, D**). Similarly, beads activated upstream kinases MEK1/2 (**Fig. 5E, F**) and cRaf (**Fig. 5G, H**); however, their phosphorylation was not sustained. Thus, in contrast to the sustained activation of MAPK/ERK that we saw during amastigote uptake, beads had only a transient effect on ERK1/2 activation.

### 3.6 Trametinib increases the intensity of actin labeled in phagocytic cups

Next, we explored the mechanisms by which trametinib affected *Leishmania* amastigote internalization. We first characterized trametinib’s effects on phagocytic cup formation by performing an internalization assay with amastigotes as described above, with the addition of fluorescent phalloidin to label phagocytic cups. We found that phalloidin labeling around internalizing parasites was significantly brighter in trametinib-treated Mϕs on average than DMSO-treated Mϕs, suggesting direct effects on actin polymerization during the formation of actin-rich phagocytic cups (**Supplemental Fig. 4A-C**). We next calculated the percentages of Mϕs infected by amastigotes and the average numbers of amastigotes per infected Mϕ. We found that both were affected by trametinib at approximately the values expected from the decrease in phagocytic indexes (**Supplemental Fig. 4D-E**).

### 3.7 Trametinib decreases lesion size and parasite burden in a murine model of cutaneous leishmaniasis

To test whether the MAPK/ERK pathway is necessary for *Leishmania* pathogenesis *in vivo*, we employed a C57BL/6 mouse model of cutaneous leishmaniasis using *L. amazonensis.* Footpad swelling (*i.e*., lesion size) was measured in infected mice treated with 2 mg/kg of body weight/day of oral trametinib or drug vehicle (control) starting 7 days (d) before inoculation and continuing for the experimental duration of 14-16 weeks (w) [16, 17] (**Fig. 6A**). Compared to controls, trametinib-treated mice developed smaller lesions (**Fig. 6B**), which contained fewer parasites in lesions (**Fig. 6C**). The decreased number of parasites found within lesions may help to explain these changes in lesion size.

Overall, in leishmaniasis, as a simplified paradigm, a Th1 response is thought to be beneficial to the host, while a Th2 response is typically considered to be deleterious. To determine whether the immunological effects of MEK1/2 inhibition were associated with the healing seen in trametinib-treated mice, we isolated draining lymph nodes from infected DMSO and trametinib-treated mice. We then profiled cytokine and chemokine secretion after stimulation with high (1 × 10^6^) or low (1 × 10^5^) doses of *L. amazonensis* lysates compared to unstimulated. A mild shift in the overall ratio of Th1 to Th2 cytokines (Th1/Th2) towards a Th2 phenotype in trametinib-treated mice was seen (**Table 1**). Notably, lymph nodes extracted from trametinib-treated mice released lower quantities of nearly all cytokines in response to *L. amazonensis* than those isolated from diluent-treated animals (**Table 2**). Trametinib-treated mice also demonstrated decreased cytokine production in response to concanavalin A (Con A) stimulation. In addition, we assessed chemokine secretion following MEK1/2 inhibition. The production of several chemokines implicated in *Leishmania* pathogenesis, including MIP-1α (CCL3) and MCP-1 (CCL2), were significantly decreased with trametinib (**Table 2**), consistent with the decrease in cytokines/chemokines overall.

**Table 1.** Ongoing trametinib treatment during *L. amazonensis* infection has limited effects on Th1 vs Th2 responses. Effects of trametinib during *L. amazonensis* infection on cytokine secretion. Shown are profiles of draining lymph nodes isolated from diluent (DMSO) versus trametinib-treated mice treated prior to inoculation as described in methods. High refers to high stimulation with a lysate of 1 × 10^6^ parasites and low refers to low stimulation with 1 × 10^5^ parasites. Ratios of IFNγ: IL-4, IFNγ: IL-13, and IFNγ: IL-4+10+13 were calculated from the data shown in Table 2. Trametinib may skew towards a Th2 response, which should not promote lesion healing.

| Experiment | Ratio | DMSO -<br>High | Trametinib<br>- High | DMSO -<br>Low | Trametinib -<br>Low |
| --- | --- | --- | --- | --- | --- |
| Trametinib<br>prior to<br>infection | **IFN Gamma/IL-4** | 1.86 | 1.35 | 1.75 | 1.05 |
|  | **IFN Gamma/IL-13** | 0.78 | 0.11 | 0.75 | 0.09 |
|  | **IFN Gamma/IL-4+**<br>**IL-10+ IL-13** | 0.55 | 0.10 | 0.52 | 0.08 |

**Table 2.** Cytokine and chemokine profiles in diluent versus trametinib-treated, *Leishmania*-infected mice. Shown are profiles of draining lymph nodes isolated from DMSO versus trametinib mice treated prior to infection as described in methods. For high stimulation categories, a lysate containing 1 × 10^6^ *L. amazonensis* was added, and for low stimulation categories, a lysate containing 1 × 10^5^ parasites was added. Post 14-16 weeks, mice were sacrificed and multiplex chemokine ELISAs were performed on harvested lymph node supernatants. *P* values were determined by two-tailed *t*-tests and are listed if *P* < 0.1. Bold: *P* ≤ 0.05.

| Experiment | Cytokine | Treatment | Cytokine secretion (pg) after |  |  |  |  |  |
| --- | --- | --- | --- | --- | --- | --- | --- | --- |
|  |  |  | Con A stimulation |  | High stimulation |  | Low treatment |  |
|  |  |  | AVG | STD | AVG | STD | AVG | STD |
| Trametinib prior to infection | **IFN Gamma** | DMSO | 6.70 | 6.83 | 6.06 | 5.82 | 5.83 | 5.16 |
|  |  | Trametinib | 0.48 | 0.64 | 0.73 | 1.05 | 0.58 | 0.75 |
|  |  | p-value | **0.01** |  | **0.01** |  | **0.005** |  |
|  | **IL-1 BETA** | DMSO | 0.88 | 0.51 | 0.97 | 0.48 | 0.73 | 0.46 |
|  |  | Trametinib | 0.45 | 0.12 | 0.42 | 0.11 | 0.39 | 0.09 |
|  |  | p-value | **0.02** |  | **0.002** |  | **0.03** |  |
|  | **IL-12/IL23p40** | DMSO | 5.56 | 2.97 | 5.24 | 2.49 | 6.04 | 3.06 |
|  |  | Trametinib | 3.28 | 1.18 | 3.39 | 0.79 | 3.53 | 1.31 |
|  |  | p-value | **0.04** |  | **0.04** |  | **0.03** |  |
|  | **IL-13** | DMSO | 5.08 | 3.04 | 7.72 | 2.47 | 7.78 | 3.07 |
|  |  | Trametinib | 7.01 | 2.27 | 6.83 | 1.10 | 6.57 | 1.15 |
|  |  | p-value | NS |  | NS |  | NS |  |
|  | **IL-17A** | DMSO | 0.48 | 0.24 | 0.65 | 0.14 | 0.53 | 0.27 |
|  |  | Trametinib | 0.59 | 0.09 | 0.74 | 0.25 | 0.53 | 0.25 |
|  |  | p-value | NS |  | NS |  | NS |  |
|  | **IL-4** | DMSO | 2.36 | 1.26 | 3.26 | 1.89 | 3.32 | 2.57 |
|  |  | Trametinib | 0.49 | 0.64 | 0.54 | 0.68 | 0.55 | 0.70 |
|  |  | p-value | **0.0006** |  | **0.0004** |  | **0.004** |  |
|  | **TNF ALPHA** | DMSO | 2.98 | 1.6 | 2.68 | 2.10 | 2.45 | 1.22 |
|  |  | Trametinib | 2.85 | 2.30 | 2.23 | 0.96 | 1.75 | 1.45 |
|  |  | p-value | NS |  | NS |  | NS |  |
|  | **IL-10** | DMSO | .008 | .007 | .02 | .009 | .009 | .01 |
|  |  | Trametinib | 8.94 | 8.78 | 8.36 | 10.44 | 5.16 | 1.06 |
|  |  | p-value | **0.005** |  | **0.02** |  | **0.0001** |  |
|  | **MCP-1** | DMSO | 25.56 | 14.87 | 25.07 | 12.31 | 27.9<br>0 | 10.54 |
|  |  | Trametinib | 7.23 | 6.43 | 7.23 | 5.87 | 7.23 | 6.56 |
|  |  | p-value | **0.002** |  | **0.0006** |  | **0.0001** |  |
|  | **MIP-1 ALPHA** | DMSO | 4.69 | 3.95 | 3.93 | 2.77 | 4.62 | 2.96 |
|  |  | Trametinib | 1.36 | 0.41 | 1.44 | 0.86 | 1.44 | 0.88 |
|  |  | p-value | **0.02** |  | **0.01** |  | **0.004** |  |

We further tested whether MEK1/2 inhibition with trametinib reduces pathogenesis after the development of symptoms by infecting mice as above. However, we started diluent vs trametinib at 6 w after inoculation (*i.e.*, once lesions were clearly visible) and continued therapy for the remaining experimental duration (**Fig. 6D**). Even with this delayed treatment, trametinib-treated mice reproducibly had smaller lesions (**Fig. 6E**). They also had significantly lower parasite burdens than those of controls (**Fig. 6F**), which at least in part may explain these results. We also saw similar immunological responses in mice treated with trametinib initiated partway through the experiment as we did for the mice treated prior to *Leishmania* infection (**Supplemental Table 1, Supplemental Table 2**) [53, 54]. Overall, our findings demonstrate that trametinib administration limits the manifestations of leishmaniasis in a mouse model, even if it is started after lesions have developed.

## 4. DISCUSSION

Here, we have shown that efficient *Leishmania* uptake by Mϕs and infection in mice requires MAPK/ERK. While previous studies have shown that ERK1/2 activation post-infection by *Leishmania* alters various host signaling pathways, leading to increased intracellular survival and replication of these parasites [36, 55], our results establish an earlier role for host ERK1/2 activity during the *Leishmania* life cycle, and show that ERK1/2 is necessary for efficient *Leishmania* uptake into Mϕs. Furthermore, our results delineate new activators of ERK signaling, and suggest that MAPK/ERK affects actin polymerization. In combination with prior data, we propose a model whereby, following engagement of the FcγR, host Arg/Abl and SYK stimulate Raf, MEK, and ERK1/2 phosphorylation, leading to amastigote uptake (**Fig. 6G**). Moreover, *in vivo* inhibition of MEK1/2 by trametinib in a mouse model of cutaneous leishmaniasis significantly limits lesion development and parasite burden, even when trametinib is started after these lesions have developed. Our results highlight a critical role for MAPK/ERK signaling in *Leishmania* uptake and further previously published studies of its role in the pathogenesis of leishmaniasis.

Our data indicate that Arg/Abl and SYK are required for amastigote-mediated ERK1/2 phosphorylation following FcγR activation, which has not been demonstrated previously. Our prior studies of host signaling required for *Leishmania* uptake revealed that amastigotes bind directly to FcγR [15] and activate SFK-Arg pathways [16] to induce uptake. We conclude from our combined results that an FcγR-SFK-Arg-SYK signaling pathway functions upstream of and activates MAPK/ERK family kinases, which in turn affects the amount of polymerized actin in phagocytic cups. We previously published similar effects on actin polymerization using the SYK inhibitor entospletinib during *Leishmania* uptake [18]. Interestingly, pharmacological inhibition of MEK1/2 and ERK1/2 in RAW 264.7 cells significantly reduce, but does not completely abolish, *Leishmania* uptake. One explanation for these results is that kinase inhibition via these compounds may be incomplete. Another is that *Leishmania* uptake may be mediated via additional host signaling pathways. Supporting the latter hypothesis, we find that low or sub-EC_50_ doses of PMA, which activates ERK1/2 through an SFK-Arg-SYK-independent manner, rescues trametinib inhibition of *Leishmania* amastigote uptake in RAW 264.7 cells, as well as in *SYK^−/−^*and *Arg^−/−^Abl^−/−^* BMDM. PMA is a known activator of PKC [26, 56-60]. However, we find that PKC activity is only slightly increased during amastigote infection of WT/DMSO-treated Mϕs, suggesting that PKC may play only a minor role during typical amastigote uptake. Ongoing work in the laboratory is directed at uncovering additional novel uptake mechanisms for *Leishmania*. In addition, although drug combinations typically are not used to treat leishmaniasis, they are the standard of care for many other infections, such as HIV or tuberculosis. Combination therapy with multiple kinase inhibitors to inhibit several uptake modulators at once may provide additional benefit for the treatment of leishmaniasis over single host cell kinase inhibitors alone.

Our data using MEK/ERK inhibitors indicate that there are differences in MAPK/ERK signaling within Mϕs depending on whether *Leishmania* amastigotes or IgG-coated beads are undergoing internalization. These results differ from most available literature that concludes that *Leishmania* primarily hijacks ongoing phagocytic pathways to facilitate its uptake. We instead suggest that there may be alternative, perhaps partly redundant, pathways that may support the uptake of *Leishmania* over the phagocytosis of beads, and vice versa. Our future experiments will continue to compare and contrast the signaling pathways required for these different uptake mechanisms between opsonized beads and *Leishmania* parasites.

Our experiments using a mouse model for *Leishmania* infection, in combination with prior studies [33], show that MAPK/ERK are required for maximal *Leishmania* pathogenesis during cutaneous leishmaniasis. One highly plausible explanation for the decreased lesion size and parasite burden that we see in trametinib-treated mice is our previously documented inhibition of *Leishmania* uptake, in combination with previous studies suggesting MAPK/ERK signaling affects parasite survival in Mϕs [26–33]. Consistent with this hypothesis, fewer parasites are found in lesions from trametinib-treated mice than in control mice. Alternately, lesion size variations after *L. amazonensis* infection might result from known trametinib limitation of host inflammatory and immunological responses. Many pathogens hinder immune responses by targeting host intracellular signaling pathways, including MAPK pathways [59, 61]; and MAPKs have been shown to play roles in the immune response to leishmaniasis [26–33]. The murine immune response to leishmaniasis is complicated and varies depending on the species of *Leishmania* and the mouse strain used. As a simplified paradigm, Th1 responses are generally protective, but Th2 responses are harmful to the host [28, 31, 32]. We find that all cytokines assessed following *Leishmania* antigen stimulation in lymph nodes isolated from infected trametinib-treated mice are reduced from the levels seen in diluent-treated mice, similar to our prior results using bosutinib [16]. One possible explanation is that fewer cytokines are generated because the lesions contain fewer parasites. Another is that trametinib could instead dampen pathways important for maintaining an active inflammatory response to the ongoing host response to infection. Supporting this second possibility, cytokines measured after Con A stimulation (an antigen-independent mitogen used as an alternative T cell stimulus) are significantly reduced by MEK1/2 inhibition. Overall, the moderate shift in the total ratio of Th1 to Th2 cytokines (Th1/Th2) toward a Th2 phenotype that we see theoretically should exacerbate disease rather than alleviate it. Reduced levels of proinflammatory chemokines like MIP-1 (CCL3) and MCP-1 (CCL2), on the other hand, may be responsible for reducing lesion size by restricting the number of immune cells, especially phagocytic cells, which reach the site of infection. We do note a surprisingly strong IL-10 response in the trametinib-treated mice, considering that ERK activation has been shown to be required for IL-10 production by *Leishmania*-infected Mϕs [62]. However, others have found that trametinib treatment increases IL-10 production in mouse tissues [63]. Further work is needed to delineate the regulation of immune responses to *Leishmania* infection and the resulting effects on lesion worsening versus healing.

Finally, and of particular interest, we have demonstrated that there are reductions in lesion size and parasite burden if trametinib is used in a *L. amazonensis* mouse model of cutaneous leishmaniasis, even if it is started after foot swelling has developed. Compared to our previous results, the data shown here strongly suggest that trametinib is more effective for reducing lesion size than imatinib [17] or bosutinib [16], which target signaling molecules in the same pathway. Trametinib is used clinically to treat several cancers, making it feasible that this drug could be repurposed to treat an infectious disease, and facilitating potential clinical trials for its use against leishmaniasis. Our data imply that inhibiting a signaling pathway downstream of SFK-Arg-SYK might improve results over inhibiting kinases upstream in the pathway. Our results also give credence to our overarching hypothesis that targeting host cell pathways, particularly ones that are required for multiple aspects of the parasite life cycle and pathogenesis, may permit treatment of an infectious disease such as leishmaniasis.

## 5. CONCLUSIONS

In summary, we have demonstrated that Arg-SYK signaling through MAPK/ERK facilitates *Leishmania* uptake by Mϕ. Furthermore, ERK activity helps govern the subsequent pathogenesis and disease progression of leishmaniasis in the mouse model. Due to the relative specificity of newer-generation MAPK/ERK small molecule inhibitors like trametinib, it is possible to selectively target MEK1/2 and ERK1/2 for potential therapeutic intervention. Our findings demonstrate the significance of MAPK/ERK signaling during *Leishmania* entry into phagocytic cells. Our future studies will investigate the use of host-cell-active drugs, potentially in conjunction with other kinase inhibitors or existing antiparasitics, to improve efficacy, decrease toxicity, and limit the likelihood of developing resistance. Although trametinib is not completely without side effects, this combination strategy could be used to mitigate any risk of toxicity arising due to targeting essential signaling pathways. Furthermore, our results support the novel possibility that medications that target conserved host cell processes rather than infectious pathogens might be employed to treat a wide spectrum of intracellular pathogens.

## Supporting information

Bioluminescence example - other supplemental material

## Abbreviations

BCA: Bicinchoninic acid
BMDM: bone marrow-derived macrophages
CC: cytotoxic concentration
CR3: complement receptor 3
FcγR: Fc gamma Receptor
h: hour
IC: inhibitory concentration
IL: Interleukin
INFγ: Interferon gamma
Mϕ: macrophage
m: minutes
MAPK/ERK: Mitogen-activated protein kinases/Extracellular signal regulated kinases
PKC: Protein Kinase C
PVDF: polyvinylidene difluoride
RAF: Rapidly Accelerated Fibrosarcoma
RIPA: radioimmunoprecipitation assay
p-: phosphorylated
PMA: Phorbol 12-myristate 13-acetate
SD: standard deviation
SFK: Src family kinases
SYK: spleen tyrosine kinase
TNF: Tumor Necrosis Factor

## 6. ACKNOWLEDGEMENTS

## 6.1 Overview

We thank Drs. Neal M. Alto, Melanie H. Cobb, Margaret A. Phillips, James J. Collins, and Michael L. Reese (all at UT Southwestern) for helpful discussions and feedback. Members of the UT Southwestern Departments of Pediatrics, Biochemistry, Microbiology, and Pharmacology provided valuable feedback and permitted common equipment use. Francis Khuong and Catherine Trice provided expert technical assistance and helped to edit this manuscript. Dr. Diane McMahon-Pratt (Yale University) generously donated antileishmanial antibodies. Dr. Anthony J. Koleske (Yale University) provided the *Arg^−/−^Abl^flox/flox^* mice. The UT Southwestern Microarray Core facilitated cytokine/chemokine profiling, the High Throughput Screening Core assisted with data visualization and quantification, and the Live Cell Imaging Core helped us with collecting confocal microscopy images.

## 6.4 Data and materials availability

All data are available within the main text of the manuscript or the supplemental materials. Any materials generated for this manuscript are freely available upon request.

## 9. LIST OF SUPPLEMENTARY MATERIALS

**Supplemental Figure 1.** MAPK/ERK inhibitors decrease uptake and survival of *L. amazonensis* promastigotes and amastigotes.

**Supplemental Figure 2.** Inhibiting kinases upstream of ERK during *L. amazonensis* uptake decreases ERK phosphorylation.

**Supplemental Figure 3.** Inhibiting ERK1/2 has minimal effects on the uptake of opsonized beads.

**Supplemental Figure 4.** Actin-based phagocytic cups are brighter in trametinib-treated Mϕs taking up *Leishmania* amastigotes compared to DMSO-treated Mϕs.

**Supplemental Table 1.** Trametinib initiated during *L. amazonensis* infection has limited effects on Th1 vs Th2 responses.

**Supplemental Table 2.** Cytokine and chemokine profiles from *Leishmania*-infected mice treated once lesions are visible.

**Supplemental Table 3.** Antibodies.

## REFERENCES AND NOTES

1. Gossage SM, Rogers ME, Bates PA. Two separate growth phases during the development of Leishmania in sand flies: implications for understanding the life cycle. Int J Parasitol. 2003;33(10):1027–34. doi: 10.1016/s0020-7519(03)00142-5. PubMed PMID: 13129524; PubMed Central PMCID: PMCPMC2839921.

2. Markle WH, Makhoul K. Cutaneous leishmaniasis: recognition and treatment. Am Fam Physician. 2004;69(6):1455-60. PubMed PMID: 15053410.

3. McCall LI, Zhang WW, Matlashewski G. Determinants for the development of visceral leishmaniasis disease. PLoS Pathog. 2013;9(1):e1003053. doi: 10.1371/journal.ppat.1003053. PubMed PMID: 23300451; PubMed Central PMCID: PMCPMC3536654.

4. Georgiadou SP, Makaritsis KP, Dalekos GN. Leishmaniasis revisited: Current aspects on epidemiology, diagnosis and treatment. J Transl Int Med. 2015;3(2):43–50. doi: 10.1515/jtim-2015-0002. PubMed PMID: 27847886; PubMed Central PMCID: PMCPMC4936444.

5. Steverding D. The history of leishmaniasis. Parasit Vectors. 2017;10(1):82. Epub 2017/02/17. doi: 10.1186/s13071-017-2028-5. PubMed PMID: 28202044; PubMed Central PMCID: PMCPMC5312593.

6. Gonzalez C, Wang O, Strutz SE, Gonzalez-Salazar C, Sanchez-Cordero V, Sarkar S. Climate change and risk of leishmaniasis in North America: predictions from ecological niche models of vector and reservoir species. PLoS Negl Trop Dis. 2010;4(1):e585. Epub 2010/01/26. doi: 10.1371/journal.pntd.0000585. PubMed PMID: 20098495; PubMed Central PMCID: PMCPMC2799657.

7. McIlwee BE, Weis SE, Hosler GA. Incidence of endemic human cutaneous leishmaniasis in the United States. JAMA Dermatol. 2018;154(9):1032–9. doi: 10.1001/jamadermatol.2018.2133. PubMed PMID: 30046836; PubMed Central PMCID: PMCPMC6143046.

8. Croft SL, Sundar S, Fairlamb AH. Drug resistance in leishmaniasis. Clin Microbiol Rev. 2006;19(1):111–26. Epub 2006/01/19. doi: 10.1128/CMR.19.1.111-126.2006. PubMed PMID: 16418526; PubMed Central PMCID: PMCPMC1360270.

9. Elmahallawy EK, Agil A. Treatment of leishmaniasis: a review and assessment of recent research. Curr Pharm Des. 2015;21(17):2259–75. Epub 2014/12/30. doi: 10.2174/1381612821666141231163053. PubMed PMID: 25543123.

10. Mann S, Frasca K, Scherrer S, Henao-Martinez AF, Newman S, Ramanan P, et al. A review of leishmaniasis: current knowledge and future directions. Curr Trop Med Rep. 2021;8(2):121–32. Epub 2021/03/23. doi: 10.1007/s40475-021-00232-7. PubMed PMID: 33747716; PubMed Central PMCID: PMCPMC7966913.

11. Ponte-Sucre A, Gamarro F, Dujardin JC, Barrett MP, Lopez-Velez R, Garcia- Hernandez R, et al. Drug resistance and treatment failure in leishmaniasis: A 21st century challenge. PLoS Negl Trop Dis. 2017;11(12):e0006052. Epub 2017/12/15. doi: 10.1371/journal.pntd.0006052. PubMed PMID: 29240765; PubMed Central PMCID: PMCPMC5730103.

12. Handman E, Bullen DV. Interaction of *Leishmania* with the host macrophage. Trends Parasitol. 2002;18(8):332–4. doi: 10.1016/s1471-4922(02)02352-8. PubMed PMID: 12377273.

13. Alexander J, Russell DG. The interaction of Leishmania species with macrophages. Adv Parasitol. 1992;31:175–254. Epub 1992/01/01. doi: 10.1016/s0065-308x(08)60022-6. PubMed PMID: 1496927.

14. Kima PE. The amastigote forms of Leishmania are experts at exploiting host cell processes to establish infection and persist. Int J Parasitol. 2007;37(10):1087–96. doi: 10.1016/j.ijpara.2007.04.007. PubMed PMID: 17543969; PubMed Central PMCID: PMCPMC2043126.

15. Ueno N, Wilson ME. Receptor-mediated phagocytosis of Leishmania: implications for intracellular survival. Trends Parasitol. 2012;28(8):335–44. doi: 10.1016/j.pt.2012.05.002. PubMed PMID: 22726697; PubMed Central PMCID: PMCPMC3399048.

16. Wetzel DM, Rhodes EL, Li S, McMahon-Pratt D, Koleske AJ. The Src kinases Hck, Fgr and Lyn activate Arg to facilitate IgG-mediated phagocytosis and Leishmania infection. J Cell Sci. 2016;129(16):3130–43. Epub 2016/07/01. doi: 10.1242/jcs.185595. PubMed PMID: 27358479; PubMed Central PMCID: PMCPMC5004897.

17. Wetzel DM, McMahon-Pratt D, Koleske AJ. The Abl and Arg kinases mediate distinct modes of phagocytosis and are required for maximal *Leishmania* infection. Mol Cell Biol. 2012;32(15):3176–86. Epub 2012/06/06. doi: 10.1128/MCB.00086-12. PubMed PMID: 22665498; PubMed Central PMCID: PMCPMC3434515.

18. Ullah I, Barrie U, Kernen RM, Mamula ET, Khuong FTH, Booshehri LM, et al. Src- and Abl-family kinases activate spleen tyrosine kinase to maximize phagocytosis and *Leishmania* infection. J Cell Sci. 2023;136(14). Epub 2023/06/26. doi: 10.1242/jcs.260809. PubMed PMID: 37357611; PubMed Central PMCID: PMCPMC10399977.

19. Greuber EK, Pendergast AM. Abl family kinases regulate FcgammaR-mediated phagocytosis in murine macrophages. J Immunol. 2012;189(11):5382–92. Epub 2012/10/27. doi: 10.4049/jimmunol.1200974. PubMed PMID: 23100514; PubMed Central PMCID: PMCPMC3504141.

20. Ortiz MA, Mikhailova T, Li X, Porter BA, Bah A, Kotula L. Src family kinases, adaptor proteins and the actin cytoskeleton in epithelial-to-mesenchymal transition. Cell Commun Signal. 2021;19(1):67. Epub 2021/07/02. doi: 10.1186/s12964-021-00750-x. PubMed PMID: 34193161; PubMed Central PMCID: PMCPMC8247114.

21. Cortez D, Reuther G, Pendergast AM. The Bcr-Abl tyrosine kinase activates mitogenic signaling pathways and stimulates G1-to-S phase transition in hematopoietic cells. Oncogene. 1997;15(19):2333–42. Epub 1997/12/11. doi: 10.1038/sj.onc.1201400. PubMed PMID: 9393877.

22. Jin A, Kurosu T, Tsuji K, Mizuchi D, Arai A, Fujita H, et al. BCR/ABL and IL-3 activate Rap1 to stimulate the B-Raf/MEK/Erk and Akt signaling pathways and to regulate proliferation, apoptosis, and adhesion. Oncogene. 2006;25(31):4332–40. Epub 2006/03/07. doi: 10.1038/sj.onc.1209459. PubMed PMID: 16518411.

23. Yang S, Liu G. Targeting the Ras/Raf/MEK/ERK pathway in hepatocellular carcinoma. Oncol Lett. 2017;13(3):1041–7. Epub 2017/04/30. doi: 10.3892/ol.2017.5557. PubMed PMID: 28454211; PubMed Central PMCID: PMCPMC5403244.

24. Guo YJ, Pan WW, Liu SB, Shen ZF, Xu Y, Hu LL. ERK/MAPK signalling pathway and tumorigenesis. Exp Ther Med. 2020;19(3):1997-2007. Epub 2020/02/28. doi: 10.3892/etm.2020.8454. PubMed PMID: 32104259; PubMed Central PMCID: PMCPMC7027163.

25. Eblen ST. Extracellular-Regulated Kinases: signaling from Ras to ERK substrates to control biological outcomes. Adv Cancer Res. 2018;138:99–142. Epub 2018/03/20. doi: 10.1016/bs.acr.2018.02.004. PubMed PMID: 29551131; PubMed Central PMCID: PMCPMC6007982.

26. Verin AD, Liu F, Bogatcheva N, Borbiev T, Hershenson MB, Wang P, et al. Role of ras-dependent ERK activation in phorbol ester-induced endothelial cell barrier dysfunction. Am J Physiol Lung Cell Mol Physiol. 2000;279(2):L360–70. Epub 2000/08/05. doi: 10.1152/ajplung.2000.279.2.L360. PubMed PMID: 10926560.

27. Wu X, Dayanand KK, Thylur RP, Norbury CC, Gowda DC. Small molecule-based inhibition of MEK1/2 proteins dampens inflammatory responses to malaria, reduces parasite load, and mitigates pathogenic outcomes. J Biol Chem. 2017;292(33):13615–34. Epub 2017/07/07. doi: 10.1074/jbc.M116.770313. PubMed PMID: 28679535; PubMed Central PMCID: PMCPMC5566520.

28. Martinez PA, Petersen CA. Chronic infection by *Leishmania amazonensis* mediated through MAPK ERK mechanisms. Immunol Res. 2014;59(1-3):153–65. Epub 2014/05/20. doi: 10.1007/s12026-014-8535-y. PubMed PMID: 24838145; PubMed Central PMCID: PMCPMC4686340.

29. Shadab M, Ali N. Evasion of host defence by *Leishmania donovani*: subversion of signaling pathways. Mol Biol Int. 2011;2011:343961. Epub 2011/11/18. doi: 10.4061/2011/343961. PubMed PMID: 22091401; PubMed Central PMCID: PMCPMC3199940.

30. Olivier M, Gregory DJ, Forget G. Subversion mechanisms by which Leishmania parasites can escape the host immune response: a signaling point of view. Clin Microbiol Rev. 2005;18(2):293–305. Epub 2005/04/16. doi: 10.1128/CMR.18.2.293-305.2005. PubMed PMID: 15831826; PubMed Central PMCID: PMCPMC1082797.

31. Ghalib HW, Whittle JA, Kubin M, Hashim FA, el-Hassan AM, Grabstein KH, et al. IL-12 enhances Th1-type responses in human *Leishmania donovani* infections. J Immunol. 1995;154(9):4623-9. Epub 1995/05/01. PubMed PMID: 7722314.

32. Bhardwaj S, Srivastava N, Sudan R, Saha B. Leishmania interferes with host cell signaling to devise a survival strategy. J Biomed Biotechnol. 2010;2010:109189. Epub 2010/04/17. doi: 10.1155/2010/109189. PubMed PMID: 20396387; PubMed Central PMCID: PMCPMC2852600.

33. Yang Z, Mosser DM, Zhang X. Activation of the MAPK, ERK, following *Leishmania amazonensis* infection of macrophages. J Immunol. 2007;178(2):1077–85. doi: 10.4049/jimmunol.178.2.1077. PubMed PMID: 17202371; PubMed Central PMCID: PMCPMC2643020.

34. Mukbel RM, Patten C, Jr., Gibson K, Ghosh M, Petersen C, Jones DE. Macrophage killing of *Leishmania amazonensis* amastigotes requires both nitric oxide and superoxide. Am J Trop Med Hyg. 2007;76(4):669–75. Epub 2007/04/12. PubMed PMID: 17426168.

35. Calegari-Silva TC, Pereira RM, De-Melo LD, Saraiva EM, Soares DC, Bellio M, et al. NF-kappaB-mediated repression of iNOS expression in *Leishmania amazonensis* macrophage infection. Immunol Lett. 2009;127(1):19–26. Epub 2009/08/29. doi: 10.1016/j.imlet.2009.08.009. PubMed PMID: 19712696.

36. Boggiatto PM, Martinez PA, Pullikuth A, Jones DE, Bellaire B, Catling A, et al. Targeted extracellular signal-regulated kinase activation mediated by *Leishmania amazonensis* requires MP1 scaffold. Microbes Infect. 2014;16(4):328–36. doi: 10.1016/j.micinf.2013.12.006. PubMed PMID: 24463270; PubMed Central PMCID: PMCPMC4023638.

37. Uphoff CC, Drexler HG. Eradication of mycoplasma contaminations. Methods Mol Biol. 2013;946:15–26. Epub 2012/11/28. doi: 10.1007/978-1-62703-128-8_2. PubMed PMID: 23179823.

38. Ullah I, Gahalawat S, Booshehri LM, Niederstrasser H, Majumdar S, Leija C, et al. An antiparasitic compound from the Medicines for Malaria Venture Pathogen Box promotes *Leishmania* tubulin polymerization. ACS Infect Dis. 2020;6(8):2057–72. Epub 2020/07/21. doi: 10.1021/acsinfecdis.0c00122. PubMed PMID: 32686409.

39. Wetzel DM, Hakansson S, Hu K, Roos D, Sibley LD. Actin filament polymerization regulates gliding motility by apicomplexan parasites. Mol Biol Cell. 2003;14(2):396–406. doi: 10.1091/mbc.E02-08-0458. PubMed PMID: 12589042; PubMed Central PMCID: PMC149980.

40. Pan AA, McMahon-Pratt D. Monoclonal antibodies specific for the amastigote stage of *Leishmania pifanoi*. I. Characterization of antigens associated with stage- and species-specific determinants. J Immunol. 1988;140(7):2406–14. PubMed PMID: 2450920.

41. Champsi J, McMahon-Pratt D. Membrane glycoprotein M-2 protects against *Leishmania amazonensis* infection. Infect Immun. 1988;56(12):3272–9. PubMed PMID: 3182080; PubMed Central PMCID: PMCPMC259734.

42. Soong L, Chang CH, Sun J, Longley BJ, Jr., Ruddle NH, Flavell RA, et al. Role of CD4+ T cells in pathogenesis associated with *Leishmania amazonensis* infection. J Immunol. 1997;158(11):5374–83. PubMed PMID: 9164958.

43. Soong L, Henard CA, Melby PC. Immunopathogenesis of non-healing American cutaneous leishmaniasis and progressive visceral leishmaniasis. Semin Immunopathol. 2012;34(6):735–51. Epub 2012/10/12. doi: 10.1007/s00281-012-0350-8. PubMed PMID: 23053396; PubMed Central PMCID: PMCPMC4111229.

44. Navas A, Vargas DA, Freudzon M, McMahon-Pratt D, Saravia NG, Gomez MA. Chronicity of dermal leishmaniasis caused by *Leishmania panamensis* is associated with parasite-mediated induction of chemokine gene expression. Infect Immun. 2014;82(7):2872–80. doi: 10.1128/IAI.01133-13. PubMed PMID: 24752514; PubMed Central PMCID: PMCPMC4097649.

45. Gonzalez-Fajardo L, Fernandez OL, McMahon-Pratt D, Saravia NG. *Ex vivo* host and parasite response to antileishmanial drugs and immunomodulators. PLoS Negl Trop Dis. 2015;9(5):e0003820. Epub 2015/05/30. doi: 10.1371/journal.pntd.0003820. PubMed PMID: 26024228; PubMed Central PMCID: PMC4449175.

46. Falchook GS, Lewis KD, Infante JR, Gordon MS, Vogelzang NJ, DeMarini DJ, et al. Activity of the oral MEK inhibitor trametinib in patients with advanced melanoma: a phase 1 dose-escalation trial. Lancet Oncol. 2012;13(8):782–9. Epub 2012/07/19. doi: 10.1016/S1470-2045(12)70269-3. PubMed PMID: 22805292; PubMed Central PMCID: PMCPMC4109286.

47. Cheng Y, Tian H. Current development status of MEK inhibitors. Molecules. 2017;22(10). Epub 2017/09/29. doi: 10.3390/molecules22101551. PubMed PMID: 28954413; PubMed Central PMCID: PMCPMC6151813.

48. Wong DJ, Robert L, Atefi MS, Lassen A, Avarappatt G, Cerniglia M, et al. Antitumor activity of the ERK inhibitor SCH772984 [corrected] against BRAF mutant, NRAS mutant and wild-type melanoma. Mol Cancer. 2014;13:194. Epub 2014/08/22. doi: 10.1186/1476-4598-13-194. PubMed PMID: 25142146; PubMed Central PMCID: PMCPMC4155088.

49. Ciuffreda L, Del Bufalo D, Desideri M, Di Sanza C, Stoppacciaro A, Ricciardi MR, et al. Growth-inhibitory and antiangiogenic activity of the MEK inhibitor PD0325901 in malignant melanoma with or without BRAF mutations. Neoplasia. 2009;11(8):720–31. Epub 2009/08/04. doi: 10.1593/neo.09398. PubMed PMID: 19649202; PubMed Central PMCID: PMCPMC2713590.

50. Sun Y, Liu WZ, Liu T, Feng X, Yang N, Zhou HF. Signaling pathway of MAPK/ERK in cell proliferation, differentiation, migration, senescence and apoptosis. J Recept Signal Transduct Res. 2015;35(6):600–4. Epub 2015/06/23. doi: 10.3109/10799893.2015.1030412. PubMed PMID: 26096166.

51. Crowley MT, Costello PS, Fitzer-Attas CJ, Turner M, Meng F, Lowell C, et al. A critical role for Syk in signal transduction and phagocytosis mediated by Fcgamma receptors on macrophages. J Exp Med. 1997;186(7):1027–39. Epub 1997/10/07. doi: 10.1084/jem.186.7.1027. PubMed PMID: 9314552; PubMed Central PMCID: PMCPMC2199061.

52. Kiefer F, Brumell J, Al-Alawi N, Latour S, Cheng A, Veillette A, et al. The Syk protein tyrosine kinase is essential for Fcgamma receptor signaling in macrophages and neutrophils. Mol Cell Biol. 1998;18(7):4209–20. Epub 1998/06/25. doi: 10.1128/MCB.18.7.4209. PubMed PMID: 9632805; PubMed Central PMCID: PMCPMC109005.

53. Liu L, Mayes PA, Eastman S, Shi H, Yadavilli S, Zhang T, et al. The BRAF and MEK inhibitors dabrafenib and trametinib: effects on immune function and in combination with immunomodulatory antibodies Targeting PD-1, PD-L1, and CTLA-4. Clin Cancer Res. 2015;21(7):1639-51. Epub 2015/01/16. doi: 10.1158/1078-0432.CCR-14-2339. PubMed PMID: 25589619.

54. Shi-Lin D, Yuan X, Zhan S, Luo-Jia T, Chao-Yang T. Trametinib, a novel MEK kinase inhibitor, suppresses lipopolysaccharide-induced tumor necrosis factor (TNF)- alpha production and endotoxin shock. Biochem Biophys Res Commun. 2015;458(3):667–73. Epub 2015/02/17. doi: 10.1016/j.bbrc.2015.01.160. PubMed PMID: 25684183.

55. Boggiatto PM, Jie F, Ghosh M, Gibson-Corley KN, Ramer-Tait AE, Jones DE, et al. Altered dendritic cell phenotype in response to *Leishmania amazonensis* amastigote infection is mediated by MAP kinase, ERK. Am J Pathol. 2009;174(5):1818–26. doi: 10.2353/ajpath.2009.080905. PubMed PMID: 19349356; PubMed Central PMCID: PMCPMC2671270.

56. Cheeseman KL, Ueyama T, Michaud TM, Kashiwagi K, Wang D, Flax LA, et al. Targeting of protein kinase C-epsilon during Fcgamma receptor-dependent phagocytosis requires the epsilonC1B domain and phospholipase C-gamma1. Mol Biol Cell. 2006;17(2):799–813. Epub 2005/12/02. doi: 10.1091/mbc.e04-12-1100. PubMed PMID: 16319178; PubMed Central PMCID: PMCPMC1356590.

57. Larsen EC, Ueyama T, Brannock PM, Shirai Y, Saito N, Larsson C, et al. A role for PKC-epsilon in Fc gammaR-mediated phagocytosis by RAW 264.7 cells. J Cell Biol. 2002;159(6):939–44. Epub 2002/12/25. doi: 10.1083/jcb.200205140. PubMed PMID: 12499353; PubMed Central PMCID: PMCPMC2173999.

58. Panetti TS, Wilcox SA, Horzempa C, McKeown-Longo PJ. Alpha v beta 5 integrin receptor-mediated endocytosis of vitronectin is protein kinase C-dependent. J Biol Chem. 1995;270(31):18593–7. Epub 1995/08/04. doi: 10.1074/jbc.270.31.18593. PubMed PMID: 7543105.

59. Roy N, Chakraborty S, Paul Chowdhury B, Banerjee S, Halder K, Majumder S, et al. Regulation of PKC mediated signaling by calcium during visceral leishmaniasis. PLoS One. 2014;9(10):e110843. Epub 2014/10/21. doi: 10.1371/journal.pone.0110843. PubMed PMID: 25329062; PubMed Central PMCID: PMCPMC4201563.

60. Zheleznyak A, Brown EJ. Immunoglobulin-mediated phagocytosis by human monocytes requires protein kinase C activation. Evidence for protein kinase C translocation to phagosomes. J Biol Chem. 1992;267(17):12042–8. Epub 1992/06/15. PubMed PMID: 1376316.

61. Arthur JS, Ley SC. Mitogen-activated protein kinases in innate immunity. Nat Rev Immunol. 2013;13(9):679–92. Epub 2013/08/21. doi: 10.1038/nri3495. PubMed PMID: 23954936.

62. Buxbaum LU, Scott P. Interleukin 10- and Fcgamma receptor-deficient mice resolve *Leishmania mexicana* lesions. Infect Immun. 2005;73(4):2101–8. Epub 2005/03/24. doi: 10.1128/IAI.73.4.2101-2108.2005. PubMed PMID: 15784551; PubMed Central PMCID: PMCPMC1087424.

63. Tada S, Anazawa T, Shindo T, Yamane K, Inoguchi K, Fujimoto N, et al. The MEK inhibitor trametinib suppresses Major Histocompatibility Antigen-mismatched rejection following pancreatic islet transplantation. Transplant Direct. 2020;6(9):e591. Epub 2020/08/28. doi: 10.1097/TXD.0000000000001045. PubMed PMID: 32851124; PubMed Central PMCID: PMCPMC7423917.

