## Supplemental Material for "MAPK/ERK Activation in Macrophages Promotes *Leishmania* Internalization and Pathogenesis"

**Supplemental Figure 1**

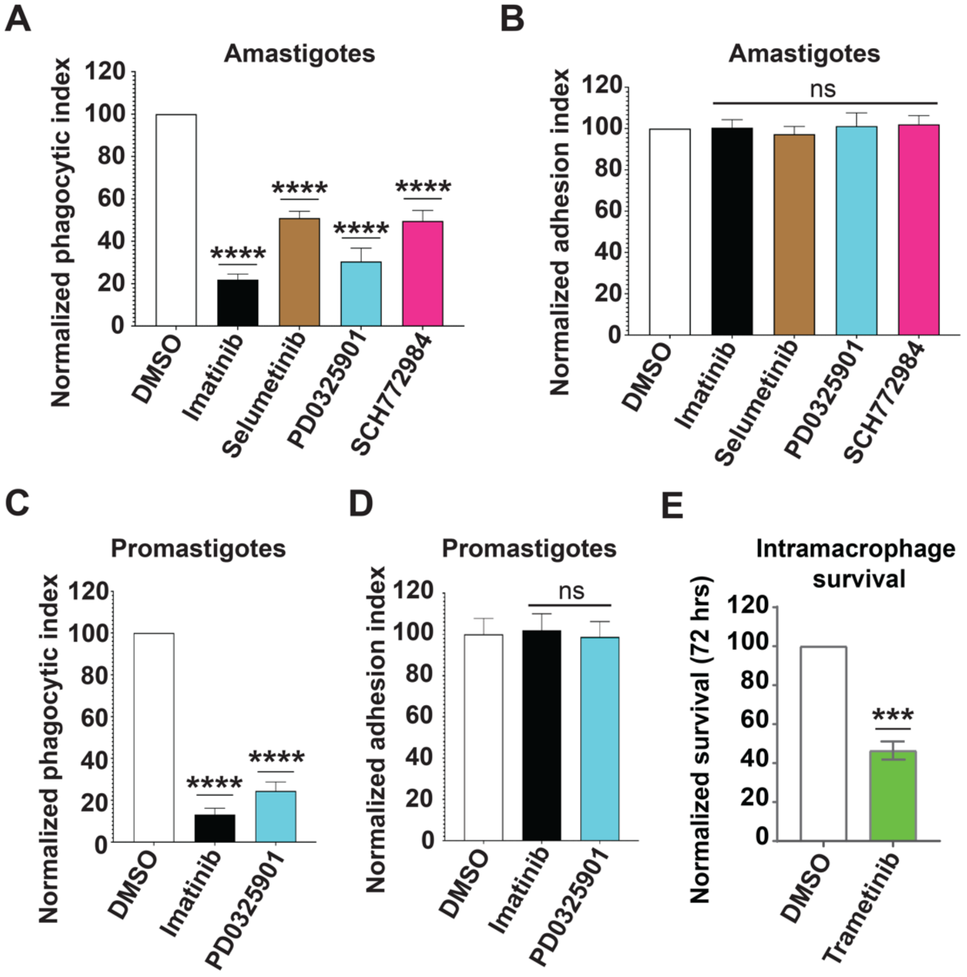


**Supplemental Figure 1. MAPK/ERK inhibitors decrease uptake and survival of *L. amazonensis* promastigotes and amastigotes.** (**A**) Selumetinib, PD0325901, and SCH772984 decrease IgG-opsonized amastigote uptake. RAW 264.7 cells were treated with 3.3 µM imatinib (Abl/Arg inhibitor), 1 µM selumetinib (MEK1/2 inhibitor), 1 µM PD0325901 (MEK1/2 inhibitor), 1 µM SCH772984 (ERK inhibitor) or DMSO and processed as in Fig 1. Graph shows the mean PI ± SD for samples, normalized to DMSO for each experiment (100%). **** $P < 0.0001$ compared to DMSO category by ANOVA (n = 6 experiments). (**B**) Imatinib, selumetinib, PD0325901, and SCH772984 do not affect adhesion. Bars show percentages of total amastigotes per 100 imatinib-treated and selumetinib-, PD0325901-, and SCH772984-treated RAW 264.7 cells relative to DMSO-treated RAW 264.7 cells. (**C**) PD0325901 decreases C3bi-promastigote uptake. RAW 264.7 cells were treated with 3.3 μM imatinib, 1 μM PD0325901, or DMSO for 2 h. 10 C3bi-opsonized promastigotes were incubated per RAW 264.7 cell and processed as for amastigote uptake. Shown is the mean PI ± SD for compound-treated samples normalized to DMSO-treated samples for each experiment (100%). **** $P < 0.0001$ compared to DMSO-treated samples by ANOVA (n = 3 experiments). (**D**) Imatinib and PD0325901 do not affect adhesion. Bars show percentages of adhered amastigotes per 100 imatinib-treated and PD0325901-treated RAW 264.7 cells relative to DMSO-treated samples from the experiment shown in panel C. **(E)** Trametinib decreases survival of intracellular amastigotes within RAW 264.7 cells. In brief, amastigotes were allowed to enter RAW 264.7 cells, and samples were washed. Samples were then incubated with trametinib or DMSO for 72 h and visualized by microscopy. Shown is the normalized number of parasites per 100 trametinib-treated RAW 264.7 cells, compared to the number of parasites per 100 DMSO-treated RAW 264.7 cells (normalized to 100% for each experiment). *** $P < 0.005$ compared to DMSO-treated samples by one sample Student's *t*-test (n = 3 experiments).

**Supplemental Figure 2**

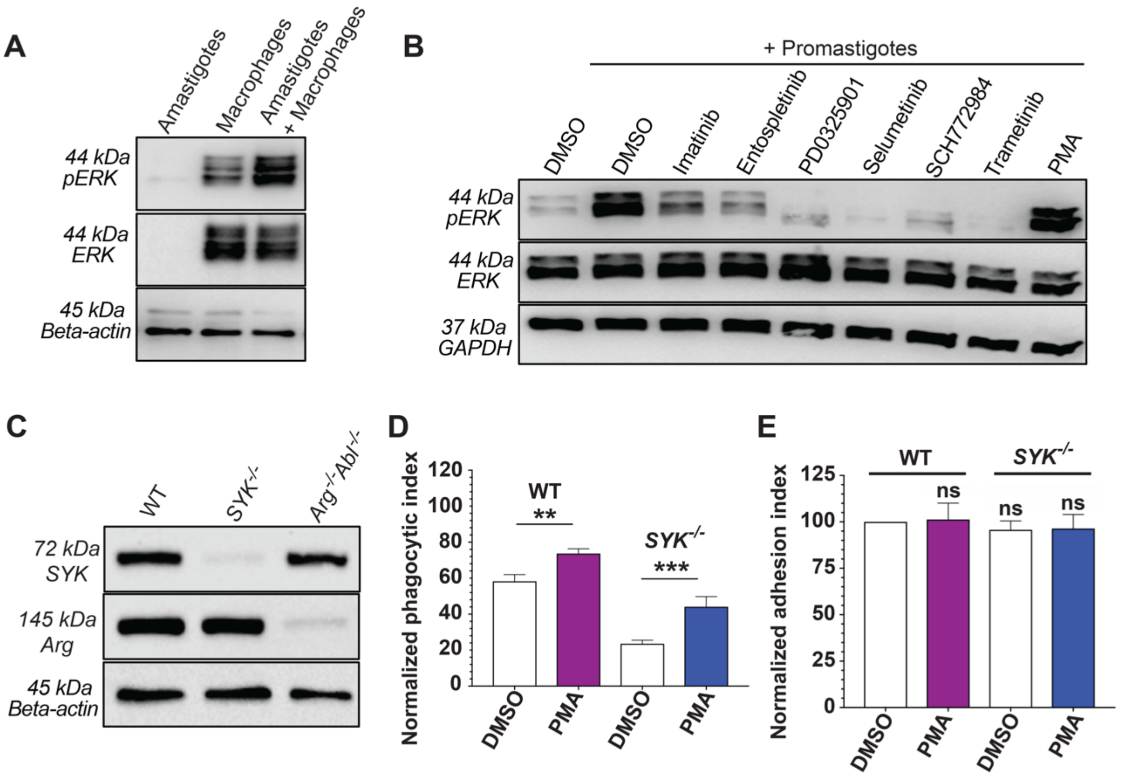


**Supplemental Figure 2. Inhibiting kinases upstream of ERK during *L. amazonensis* uptake decreases ERK phosphorylation. (A)** pERK and ERK are not detected in amastigotes by the antibodies used to perform Western blots. Equivalent total protein amounts were loaded in each lane. (**B**) Phosphorylation of ERK1/2 induced upon promastigote uptake is decreased by imatinib, entospletinib, PD0325901, selumetinib, SCH772984, and trametinib and increased by PMA. RAW 264.7 cells were treated with inhibitors or DMSO prior to adding complement-opsonized promastigotes for 30 m. Processing for immunoblotting followed. **(C)** Immunoblots of SYK and Arg expression in BMDM isolated from WT mice, *SYK*$^{flox/flox}$ LysM Cre$^{+}$ mice, (abbreviated *SYK*$^{-/-}$ when describing BMDM) or *Arg*$^{-/-}$*Abl*$^{flox/flox}$ LysM Cre$^{+}$ mice (abbreviated *Arg*$^{-/-}$*Abl*$^{-/-}$

when describing BMDM). Actin is a loading control. (**D**) Decrease in amastigote uptake seen in *SYK*$^{-/-}$ BMDM is rescued by PMA. Bars show mean ± SD of total amastigotes per 100 PMA-treated and/or *SYK*$^{-/-}$ BMDM relative to WT DMSO-treated BMDM. n = 3 biological experiments. **(E)** DMSO and PMA does not affect adhesion for the experiments shown in D.

**Supplemental Figure 3**

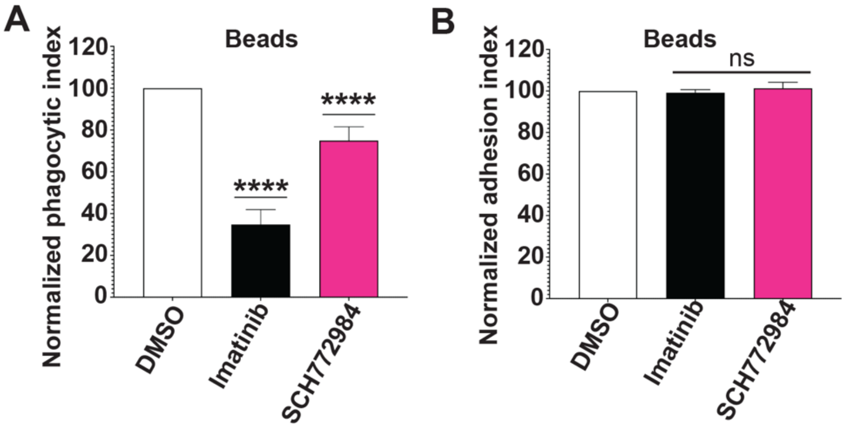


**Supplemental Figure 3. Inhibiting ERK1/2 has minimal effects on the uptake of opsonized beads.** (**A**) The ERK1/2 inhibitor SCH772984 mildly decreases phagocytosis of IgG-opsonized beads. RAW 264.7 cells were treated with 3.3 µM imatinib, 1 µM SCH772984 or DMSO, incubated with IgG-opsonized beads, and processed for immunofluorescence. Graph shows the mean phagocytic index (PI) ± standard deviation (SD) for these categories, normalized to DMSO for each experiment (100%). **** $P < 0.0001$ compared to DMSO-treated RAW 264.7 cells by ANOVA (n = 6 experiments). (**B**) Imatinib and SCH772984 do not affect adhesion of beads. Bars show percentages of total (internal + external) beads per 100 imatinib and SCH772984-treated RAW 264.7 cells relative to DMSO-treated cells from the experiments shown in panel A.

**Supplemental Figure 4**

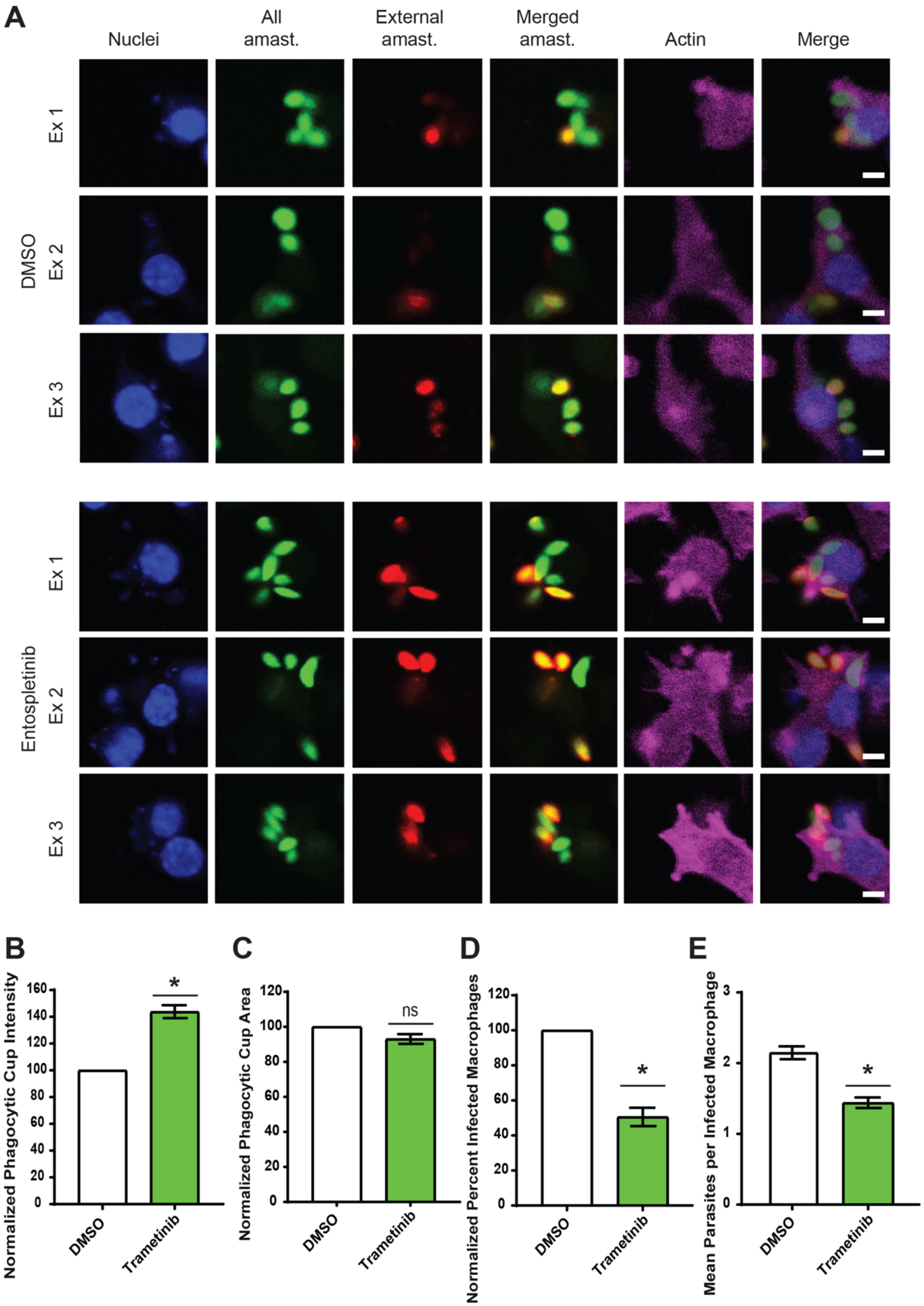

**Supplemental Figure 4. Actin-based phagocytic cups are brighter in trametinib-treated Mϕs taking up *Leishmania* amastigotes compared to DMSO-treated Mϕs.** (A) Three separate examples of phagocytic cups in DMSO versus trametinib-treated RAW 264.7 cells that are in the process of internalizing amastigotes are shown. From left to right: Mϕ nuclei (blue, stained with Hoescht); all amastigotes (green; labeled with an antibody to p8); external amastigotes (red; labeled with an antibody to p8); merged amastigotes (green/yellow/red); actin (pink; stained with Far Red 647-phalloidin); fully merged images (all colors). Scale bar = 2 μm. (**B**) Similar to what we have published for entospletinib-treated phagocytes, actin staining is brighter in trametinib-treated Mϕs than DMSO-treated Mϕs. Relative fluorescence intensity was quantified in ImageJ by an observer blinded to experimental category. Shown is the normalized mean intensity per field among at least 5 fields per category ± SE. * $P < 0.05$ by one sample, two-tailed *t*-test. (**C**) The size of phagocytic cups is similar in trametinib-treated Mϕs compared to DMSO-treated Mϕs. Relative fluorescence intensity was quantified in ImageJ by an observer blinded to experimental category and normalized to the value for DMSO-treated RAW cells. n.s., non-significant by *t*-test. (**D**) The percentages of trametinib-treated Mϕs infected by amastigotes are decreased compared to DMSO-treated Mϕs. Shown is the mean percentage of infected trametinib-treated Mϕ per category, normalized to the percentage of infected DMSO-treated Mϕs for each biological experiment, ± SE. * $P < 0.05$ by one sample, two-tailed *t*-test. (**E**) The mean number of amastigotes contained within each infected Mϕ is decreased in trametinib-treated vs DMSO-treated Mϕs, shown for each biological experiment,

± SE. * $P < 0.05$ by one sample, two-tailed *t*-test.

**Supplemental Table 1**

| Experiment | Ratio | DMSO - High | Trametinib - High | DMSO - Low | Trametinib - Low |
|---|---|---|---|---|---|
| Trametinib 6 weeks after infection | **IFN Gamma/IL-4** | 0.46 | 0.40 | 0.35 | 0.43 |
| | **IFN Gamma/IL-13** | 0.15 | 0.23 | 0.05 | 0.18 |
| | **IFN Gamma/IL-4 + IL-10+ IL-13** | 0.11 | 0.22 | 0.04 | 0.17 |

**Supplemental Table 1. Trametinib initiated during *L. amazonensis* infection has limited effects on Th1 vs Th2 responses.** Effects of trametinib treatment during *L. amazonensis* infection on cytokine secretion. Shown are profiles of draining lymph nodes isolated from DMSO versus trametinib mice treated 6 weeks after inoculation as described in methods. Ratios of IFNγ: IL-4, IFNγ: IL-13, and IFNγ:IL-4+10+13 were calculated from the data shown in Supplemental Table 2.

### Supplemental Table 2

| Experiment | Cytokine | Treatment | Cytokine secretion (pg) after | | | | | |
|---|---|---|---|---|---|---|---|---|
| | | | Con A stimulation | | High stimulation | | Low treatment | |
| | | | AVG | STD | AVG | STD | AVG | STD |
| Trametinib 6 weeks after infection | **IFN Gamma (38)** | DMSO | 3.71 | 3.07 | 11.78 | 16.42 | 7.38 | 6.07 |
| | | Trametinib | 1.46 | 1.20 | 1.80 | 1.31 | 1.64 | 1.23 |
| | | p-value | **0.04** | | 0.07 | | 0.009 | |
| | **IL-1 BETA (19)** | DMSO | 0.48 | 0.18 | 0.50 | 0.16 | 0.64 | 0.05 |
| | | Trametinib | 0.3 | 0.16 | 0.37 | 0.13 | 0.32 | 0.07 |
| | | p-value | **0.03** | | 0.06 | | **0.0001** | |
| | **IL-12/IL23p40 (64)** | DMSO | 8.98 | 3.00 | 10.80 | 4.73 | 10.50 | 4.21 |
| | | Trametinib | 2.23 | 1.09 | 2.26 | 1.15 | 2.27 | 1.14 |
| | | p-value | **0.0001** | | **0.0001** | | **0.0001** | |
| | **IL-13 (35)** | DMSO | 133.05 | 150.75 | 80.63 | 105.74 | 159.86 | 226.29 |
| | | Trametinib | 6.81 | 4.34 | 7.95 | 5.22 | 8.95 | 5.86 |
| | | p-value | **0.02** | | **0.04** | | **0.05** | |
| | **IL-17A (52)** | DMSO | 1.55 | 1.48 | 1.34 | 1.27 | 2.19 | 2.12 |
| | | Trametinib | 0.60 | 0.16 | 0.73 | 0.25 | 0.73 | 0.26 |
| | | p-value | 0.06 | | NS | | **0.04** | |
| | **IL-4 (26)** | DMSO | 11.41 | 15.15 | 25.68 | 33.79 | 21.37 | 28.79 |
| | | Trametinib | 2.79 | 2.45 | 4.53 | 4.89 | 3.79 | 3.55 |
| | | p-value | 0.09 | | 0.06 | | 0.07 | |
| | **TNF ALPHA (45)** | DMSO | 3.10 | 1.25 | 4.43 | 2.33 | 5.25 | 0.73 |
| | | Trametinib | 1.71 | 0.46 | 1.37 | 0.49 | 1.59 | 0.57 |
| | | p-value | **0.004** | | **0.0007** | | 0.0001 | |

| | | | | | | | | |
|---|---|---|---|---|---|---|---|---|
| | **IL-10 (13)** | DMSO | 3.19 | 0.35 | 8.67 | 2.47 | 15.63 | 16.16 |
| | | Trametinib | .02 | .01 | .008 | 0.005 | .009 | .006 |
| | | p-value | **0.0001** | | **0.0001** | | **0.007** | |
| | **MCP-1 (51)** | DMSO | 60.23 | 57.72 | 53.89 | 52.08 | 57.44 | 55.98 |
| | | Trametinib | 11.26 | 8.67 | 7.23 | 5.85 | 14.31 | 13.11 |
| | | p-value | **0.02** | | **0.01** | | 0.03 | |
| | **MIP-1 ALPHA (47)** | DMSO | 19.08 | 22.87 | 15.97 | 19.38 | 23.08 | 28.64 |
| | | Trametinib | 2.04 | 1.64 | 2.26 | 1.85 | 2.49 | 2.04 |
| | | p-value | **0.03** | | **0.04** | | **0.04** | |

**Supplemental Table 2. Cytokine and chemokine profiles from *Leishmania*-infected mice treated once lesions are visible.** Shown are profiles of draining lymph nodes isolated from DMSO versus trametinib mice treated 6 weeks after infection, as described in methods. Post 14-16 weeks, mice were sacrificed and multiplex chemokine ELISAs were performed on harvested lymph node supernatants. *P* values were determined by two-tailed *t*-test and are listed if $P < 0.1$. Bold $P \leq 0.05$.

### Supplemental Table 3

| Protein | Size (kDa) | Species | Company | Catalog number | Dilutions |
|---|---|---|---|---|---|
| p-ERK1/2 (p44/42 MAPK) (Thr202/Tyr204) | 44/42 | Rabbit | Cell Signaling Technologies | 9101 | 1:1000 for WB; 1:200 for IF |
| ERK1/2 (p44/42 MAPK) | 44/42 | Rabbit | Cell Signaling Technologies | 9102 | 1:1000 for WB; 1:200 for IF |
| p-c-Raf (Ser338) | 74 | Rabbit | Cell Signaling Technologies | 9427 | 1:1000 for WB |
| c-Raf | 65-75 | Rabbit | Cell Signaling Technologies | 9422 | 1:1000 for WB |
| p-MEK1/2 (Ser217/Ser221) | 45 | Rabbit | Cell Signaling Technologies | 9154 | 1:1000 for WB and IF |
| MEK1/2 | 45 | Rabbit | Cell Signaling Technologies | 9122 | 1:1000 for WB and IF |
| GAPDH (14C10) | 37 | Rabbit | Cell Signaling Technologies | 2118S | 1:1000 for WB |
| b-actin | 45 | Mouse | Cell Signaling Technologies | 3700 | 1:1000 for WB |
| p-SYK (Tyr525/526) | 72 | Rabbit | Cell Signaling Technologies | 2710T | 1:1000 for WB |
| SYK | 72 | Rabbit | Cell Signaling Technologies | 13198T | 1:500 for IF |
| SYK | 72 | Rabbit | Santa Cruz Biotechnology | sc-1240 | 1:1000 for WB |

| | | | | | |
|---|---|---|---|---|---|
| ARG | 145 | Mouse | Santa Cruz Biotechnology | sc-81154 | 1:250 for WB |
| Goat Anti-mouse IgG HRP-conjugated | - | Mouse | Cell Signaling Technologies | 7076 | 1:2000 for WB |
| Goat Anti-rabbit IgG HRP-conjugated | - | Rabbit | Cell Signaling Technologies | 7074 | 1:2000 for WB |
| Alexa Fluor 488, anti-rabbit IgG | - | Rabbit | Invitrogen | A-21206 | 1:100 for IF |
| Alexa Fluor 594, anti-rabbit IgG | - | Rabbit | Invitrogen | A-11037 | 1:100 for IF |
| Alexa Fluor 647, anti-rabbit IgG | - | Rabbit | Invitrogen | A-32733 | 1:100 for IF |
| IgG (H+L) cross-absorbed Goat anti-mouse, Alexa Fluor 488, Phalloidin | - | Mouse | Invitrogen | A-11001 | 1:250 for IF |
| IgG (H+L) cross-absorbed Goat anti-mouse, Alexa Fluor 568, Phalloidin | - | Mouse | Invitrogen | A-11004 | 1:250 for IF |
| IgG (H+L) cross-absorbed Goat anti-rabbit, Alexa Fluor 594, Phalloidin | - | Rabbit | Invitrogen | A-11012 | 1:250 for IF |
| IgG (H+L) cross-absorbed Goat anti-mouse, Alexa Fluor 647, Phalloidin | - | Mouse | Invitrogen | A-21235 | 1:250 for IF |

**Supplemental Table 3. Antibodies.** Identity of all antibodies used for western blot (WB) and immunofluorescence (IF).
