## Supplementary material for "MAPK/ERK Activation in Macrophages Promotes *Leishmania* Internalization and Pathogenesis": Bioluminescence example - other supplemental material

**Other supplemental data**

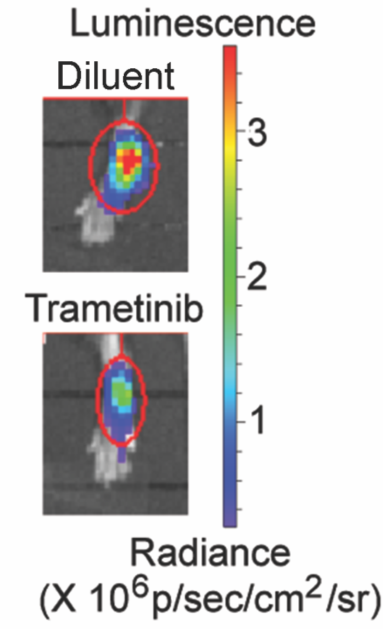


**Other supplemental data.** Bioluminescence imaging of luciferase-expressing *L. amazonensis*-infected mice treated with diluent versus trametinib. C57BL/6 mice were given 250 μl of D-luciferin intraperitoneally and imaged 8 weeks after infection/2 weeks after trametinib initiation, as in **Fig 6D-F**. This figure provides a proof-of-concept demonstration that we can detect decreases in parasite burden with bioluminescence in live mice treated with kinase inhibitors.
